# *Enterococcus faecalis* persists and replicates within epithelial cells *in vitro* and *in vivo* during wound infection

**DOI:** 10.1101/2021.09.16.460717

**Authors:** Wei Hong Tay, Ronni A.G. da Silva, Foo Kiong Ho, Kelvin K.L. Chong, Alexander Ludwig, Kimberly A. Kline

## Abstract

*Enterococcus faecalis* is a frequent opportunistic pathogen of wounds, whose infections are associated with biofilm formation, persistence, and recalcitrance toward treatment. We have previously shown that *E. faecalis* wound infection persists for at least 7 days. Here we report that viable *E. faecalis* are present within both immune and non-immune cells at the wound site up to 5 days after infection, raising the prospect that intracellular persistence contributes to chronic *E. faecalis* infection. Using an *in vitro* keratinocyte infection model, we show that a subpopulation of *E. faecalis* becomes internalized via macropinocytosis into single membrane-bound compartments, where they can survive and replicate. These intracellular *E. faecalis* can persist in late endosomes up to 72 hours after infection in the absence of colocalization with the lysosomal protease cathepsin D or apparent fusion with the lysosome, suggesting that *E. faecalis* blocks endosomal maturation. Indeed, intracellular *E. faecalis* infection results in a marked reduction in Rab7 expression, a small GTPase required for endosome-lysosome fusion. Finally, we demonstrate that intracellular *E. faecalis* derived from infected keratinocytes are significantly more efficient in reinfecting new keratinocytes. Together, these data suggest that intracellular proliferation of *E. faecalis* may contribute to its persistence in the face of a robust immune response, providing a primed reservoir of bacteria for subsequent reinfection.

**Author Summary:** *Enterococcus faecalis* is often isolated from chronic wounds. Prior to this study, *E. faecalis* has been observed within different cell types, suggesting that it can successfully colonize intracellular spaces. However, to date, little is known about the mechanisms *E. faecalis* use to survive intracellularly. Here, we describe key features of the intracellular lifestyle of *E. faecalis*. We show that *E. faecalis* exists in an intracellular state within immune cells and non-immune cells during mammalian wound infection. We show that *E. faecalis* can survive and replicate inside keratinocytes, and intracellularly replicating *E. faecalis* are primed to more efficiently cause reinfection, potentially contributing to chronic or persistent infections. In order to establish this intracellular lifestyle, *E. faecalis* is taken up by keratinocytes via macropinocytosis, whereupon it manipulates the endosomal pathway and expression of trafficking molecules required for endo-lysosomal fusion, enabling *E. faecalis* to avoid lysosomal degradation and consequent death. These results advance our understanding of *E. faecalis* pathogenesis, demonstrating mechanistically how this classic extracellular pathogen can co-opt host cells for intracellular persistence, and highlight the heterogeneity of mechanisms bacteria can use to avoid host-mediated killing in order to cause disease.

## Introduction

*Enterococcus faecalis*, traditionally considered an extracellular pathogen, is a member of the healthy human gut microbiome and a frequent opportunistic pathogen of the urinary tract and wounds. *Enterococci* are one of the most frequently isolated bacterial genera from wound infections (Dowd et al. 2008; Giacometti et al. 2000; Gjodsbol et al. 2006; Heitkamp et al. 2018; Brook and Frazier 1998); however, their pathogenic mechanisms enabling persistence in this niche are not well understood. We have previously shown in a mouse excisional wound infection model that *E. faecalis* undergo acute replication and long term persistence, leading to delayed wound healing, despite a robust innate inflammatory response at the wound site (Chong et al. 2017). These data suggest that *E. faecalis* possess mechanisms to evade the innate immune response, and indeed, we have also shown that extracellular *E. faecalis* can actively suppress NF-κB activation in macrophages (Tien et al. 2017). In addition, *E. faecalis* can persist within a variety of eukaryotic cells including macrophages (Zou and Shankar 2014; Gentry-Weeks et al. 1999; Zou and Shankar 2016), osteoblasts (Campoccia et al. 2016; Tong et al. 2016), monocytes (Baldassarri et al. 2005), endothelial cells (Millan et al. 2013), and epithelial cells (Bertuccini et al. 2002; Horsley et al. 2018; Horsley et al. 2013; Olmsted et al. 1994; Wells, Jechorek, and Erlandsen 1990; Wells et al. 1988). However, the mechanisms mediating intracellular persistence have not been reported.

In this study, we sought to understand how *E. faecalis* persist within epithelial cells and how intracellularity contributes to pathogenesis. Using a mouse model of wound infection, we found viable *E. faecalis* within both immune and non-immune cells at the wound site up to 5 days after infection. Using an *in vitro* model of keratinocyte infection, we show that *E. faecalis* is taken up into these cells via macropinocytosis, whereupon they traffic through the endocytic pathway to late endosomes, and ultimately undergo intracellular replication. Interestingly, *E. faecalis* infection results in a marked reduction of Rab7 expression, a small GTPase required for late endosome-lysosome fusion, providing a potential mechanism for intracellular survival. Finally, we show that *E. faecalis* derived from the intracellular niche are primed to more efficiently reinfect new keratinocytes. Together, our data are consistent with a model in which a subpopulation of *E. faecalis* are taken up into epithelial cells during wound infection, providing immune protection and a replicative niche, which may serve as a nidus for chronically infected wounds.

## Results

### Intracellular *E. faecalis* are present within CD45+ and CD45-cells during mouse wound infection

To determine whether *E. faecalis* persists intracellularly within infected wounds, we infected wounded mice with 10^6^ CFU of *E. faecalis* for 1, 3 and 5 days. Infected wounds were dissociated to a single cell suspension, treated with gentamicin/penicillin G to kill extracellular bacteria, immunolabeled with anti-CD45 antibody, and sorted into CD45+ immune cells and CD45-non-immune cells. These sorted cells were then lysed for the enumeration of intracellular bacteria. Consistent with literature reporting the ability of *E. faecalis* to persist within phagocytic immune cells, we recovered viable *E. faecalis* from CD45+ cells (**Figure 1A**). In addition, intracellular *E. faecalis* was also recovered from the CD45-population, up to 5 dpi (**Figure 1B**). Compared to the approximately 10^5^ CFU total recoverable *E. faecalis* (both extracellular and intracellular) within wounds at 3 and 5 dpi (Chong et al. 2017), we can estimate that approximately 1-10% of the total recovered bacterial population are intracellular at these times. These data demonstrate that *E. faecalis* can exist intracellularly during wound infection, implying it is not an exclusive extracellular pathogen.

**Figure 1.**
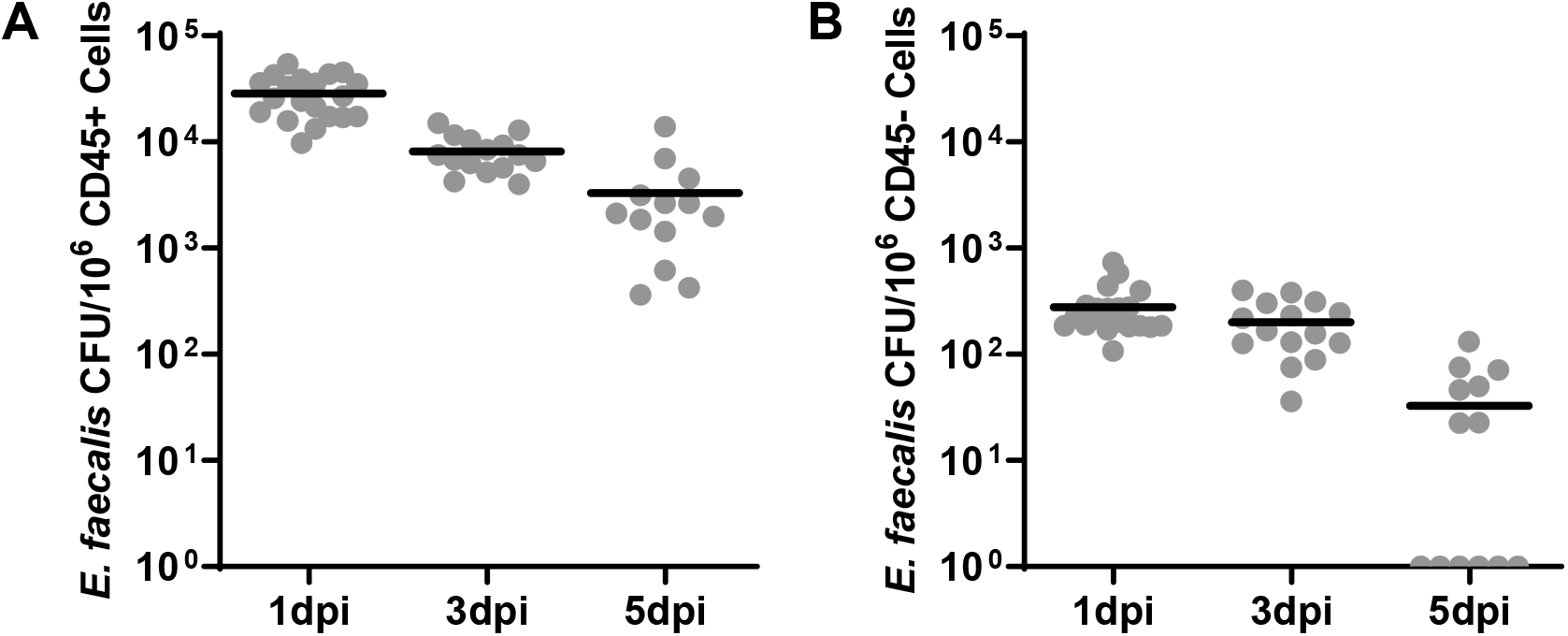
Recovery of viable *E. faecalis* within host cells during wound infection. Male C57BL/6 mice were wounded and infected with 10^6^ CFU of *E. faecalis* OG1RF. Wounds were harvested at 1, 3, or 5 days post-infection (dpi), dissociated to single cell suspension, treated with antibiotics to kill extracellular bacteria, labeled, and sorted into **(A)** CD45+ (immune) or **(B)** CD45- (non-immune) populations. CD45+ and CD45-cell populations were then lysed and plated for bacterial CFU. Each data point indicates the CFU within the sorted subpopulation from one mouse. Data shown represent at least 3 independent experiments, each of which included at least 4 mice per time point. Horizontal black lines indicate the mean for each group.

### *E. faecalis* adheres to and enters keratinocytes

To investigate the mechanisms by which *E. faecalis* infects non-immune cells at the wound site, we infected the spontaneously immortalized human keratinocyte cell line (HaCaT) with *E. faecalis* strain OG1RF at a multiplicity of infection (MOI) of 1, 10 or 100 for a period of up to 3 hours, followed by 1 hour of gentamicin/penicillin treatment to kill extracellular bacteria and quantified the intracellular bacteria. We observed that *E. faecalis* can adhere to keratinocytes at all MOI and time points **(Figure 2A)**, and intracellular *E. faecalis* were recovered as early as 1 hour post-infection (hpi) at a MOI of 10 and 100 **(Figure 2B)**. Parallel cytotoxicity experiments established that *E. faecalis* infection does not negatively affect keratinocytes at early time points of <5 hpi, even in the absence of gentamicin/penicillin (data not shown). Thus, we chose MOI 100 and no more than 3 hours of infection without antibiotics to characterize its intracellular pathogenesis. Since we recovered intracellular *E. faecalis* from infected mouse wounds in both immune and non-immune compartments, we hypothesized that *E. faecalis* could also survive within mouse fibroblasts and macrophages *in vitro*. Indeed, intracellular *E. faecalis* was recovered from RAW264.7 murine macrophages and NIH/3T3 murine fibroblasts, indicating that *E. faecalis* internalization and persistence is not cell type specific **(Supplementary Figure 1A)**. Similarly, intracellular persistence within keratinocytes was not *E. faecalis* strain specific, as the vancomycin resistant strain V583 persisted at even higher numbers within HaCaT cells **(Supplementary Figure 1B,C)**.

**Figure 2.**
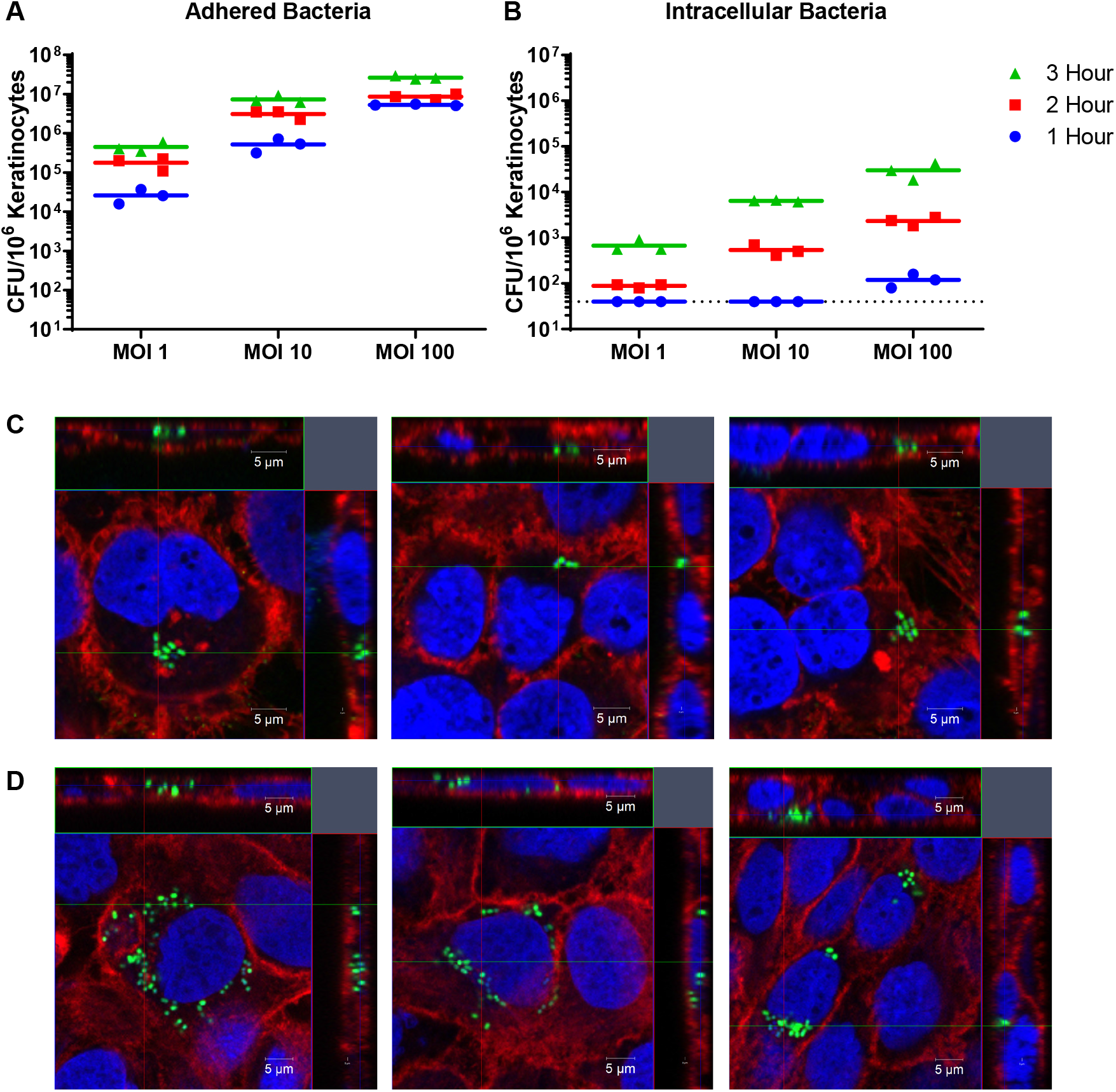
Time and dose-dependent increase of intracellular *E. faecalis* with keratinocytes *in vitro*. 10^6^ keratinocytes were infected at the indicated MOI for 1, 2, or 3 h, each followed by another 1 h of antibiotic treatment. **(A, B)** Solid lines indicate the mean CFU/well from a total of 3 independent experiments. Dashed black line indicates the limit of detection of the assay. **(C)** CLSM orthogonal view of internalized *E. faecalis* within keratinocytes at 4 hpi (3 h infection + 1 h antibiotic treatment). **(D)** CLSM orthogonal view of internalized *E. faecalis* within keratinocytes at 24 hpi (3 h infection + 21 h antibiotic treatment). **(C, D)** Blue, dsDNA stained with Hoechst 33342; green, E-GFP, *E. faecalis*; red, F-actin. Images are representative of 3 independent experiments. (See **Supplementary Figures 1 and 2** for data related to the 24 hpi time point).

To confirm the presence of *E. faecalis* within keratinocytes, we imaged keratinocytes infected with a green fluorescent protein (GFP) expressing derivative of *E. faecalis* (Debroy et al. 2012) by confocal laser scanning microscopy (CLSM). Images taken at 4 hpi, from cells infected for 3 hours followed by 1 hour gentamicin/penicillin treatment to kill extracellular bacteria, revealed 1-10 intracellular bacteria within each infected keratinocyte **(Figure 2C)**. This observation suggested either that selected infected keratinocytes can take up many *E. faecalis*, or that *E. faecalis* could replicate within these keratinocytes. To discern if *E. faecalis* can replicate within keratinocytes, we extended the period of post-infection antibiotic exposure up to 24 hpi and recovered similar, albeit slightly decreasing over time, intracellular CFU within the whole population **(Supplementary Figure 1B)**. However, within single infected keratinocytes, we visualized 10-30 *E. faecalis*, which often clustered in a perinuclear region (**Figure 2D**). At the same 24 hpi time point, we also detected clusters of fluorescent *E. faecalis* peripheral to apparently apoptotic keratinocytes (**Supplementary Figure 2**), indicative of intracellular bacteria that have either escaped from the keratinocyte or of bacteria derived from lysed keratinocytes, of which the latter may account for the slight decrease in overall intracellular CFU over time. Since we observed larger clusters of intracellular *E. faecalis* within individual keratinocytes upon extension of post-infection antibiotic exposure to 24 hpi, compared with an antibiotic exposure up to 4 hpi, these data suggest that *E. faecalis* is replicating within the cells over the subsequent 21 hrs.

**Supplementary Figure 1 (related to Figure 2).**
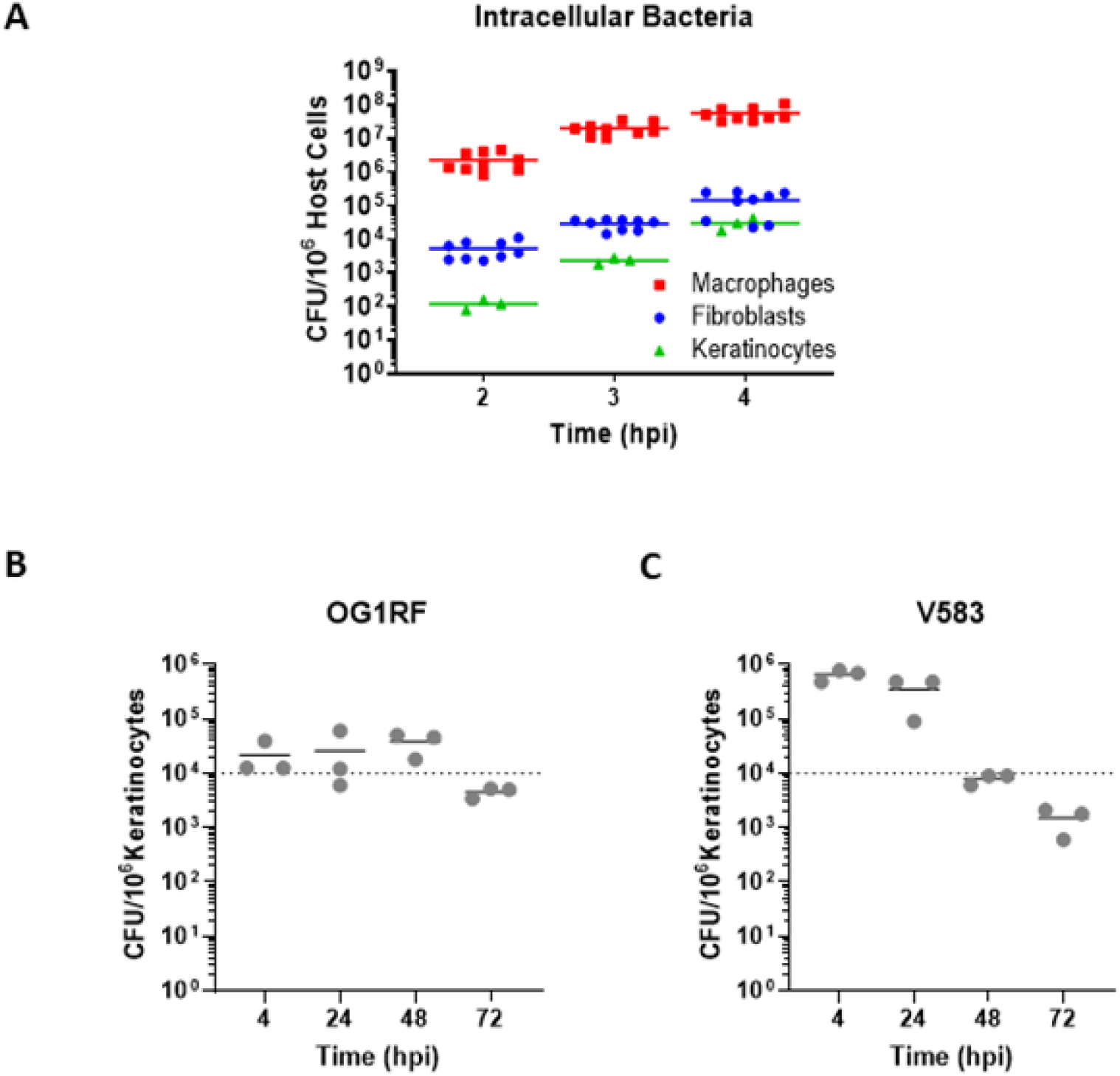
Intracellular *E. faecalis* is not cell type specific and persists for up to 72 hpi. **(A)** Solid lines indicate the mean CFU at 2-4 hpi at MOI 100 from at least 3 independent experiments. **(B and C)** HaCaTs were infected with *E. faecalis* OG1RF and V583 at MOI 100 for 3 h, followed by treatment with gentamicin/penicillin for 1, 21, 45, 69 h before lysis to obtain the intracellular population. Solid lines indicate the mean CFU from at least 2 independent experiments.

**Supplementary Figure 2 (related to Figure 2).**
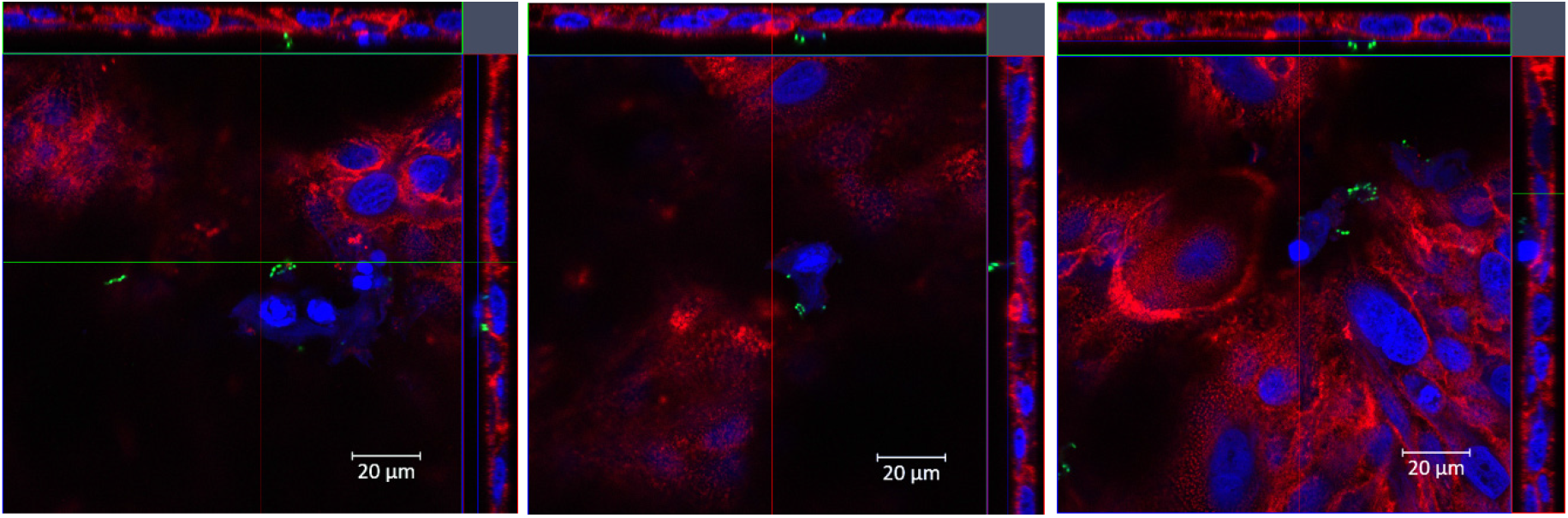
*E. faecalis* at the periphery of keratinocytes at 24 hpi. CLSM representative images of infected keratinocytes with condensed nuclei following 3 h of infection and 21 h of incubation in antibiotic laced media. Blue, dsDNA stained with Hoechst 33342; green, E-GFP *E. faecalis*; red, F-actin. Data shown are representative of at least 3 independent experiments.

### Entry of *E. faecalis* into keratinocytes is dependent on actin polymerization and PI3K signalling

Bacterial uptake into non-professional phagocytes such as epithelial cells can proceed via a number of different endogenous endocytic pathways (Haglund and Welch 2011). Previous studies have suggested that *E. faecalis* uptake into non-professional phagocytic cells is dependent on actin and microtubule polymerization, suggestive of macropinocytosis or receptor (clathrin)-mediated endocytosis (Bertuccini et al. 2002; Millan et al. 2013). To determine whether an intact cytoskeleton is important for *E. faecalis* entry into keratinocytes, we pre-treated keratinocytes with specific chemical inhibitors, prior to infection and intracellular CFU enumeration. We found that cytochalasin-D and latrunculin A, inhibitors of actin filament polymerization (Coue et al. 1987; Flanagan and Lin 1980), did not alter bacterial adhesion to keratinocytes (**Figure 3A**), but resulted in a significant 100-fold decrease in recoverable intracellular bacteria, demonstrating that actin polymerization is important for the entry process (**Figure 3B**). By contrast, colchicine, an inhibitor of microtubule polymerization (Andreu and Timasheff 1982), did not impede bacterial adhesion or uptake **(Supplementary Figure 3A)**. Since many endocytic pathways rely on an intact actin cytoskeleton (Mayor and Pagano 2007), a panel of more selective inhibitors was used to determine the mechanism of *E. faecalis* entry. Inhibitors of receptor (clathrin)- and caveolae-mediated endocytosis did not meaningfully affect bacterial adhesion or internalization, indicating that these pathways are not required for *E. faecalis* infection **(Supplementary Figure 3B,C)**. To test the role of macropinocytosis in *E. faecalis* uptake, we pre-treated keratinocytes with the phosphoinositide 3-kinase (PI3K) inhibitor wortmannin (Doherty and McMahon 2009; Wymann et al. 1996). Wortmannin did not affect *E. faecalis* adhesion to keratinocytes (**Figure 3C**), but resulted in a 10-fold decrease in intracellular CFU, as compared to the untreated controls (**Figure 3D**). Together, the dependence of *E. faecalis* internalization on actin polymerization and PI3K, and independence of receptor (clathrin)- and caveolae-mediated endocytosis, is consistent with macropinocytosis-mediated uptake.

**Figure 3.**
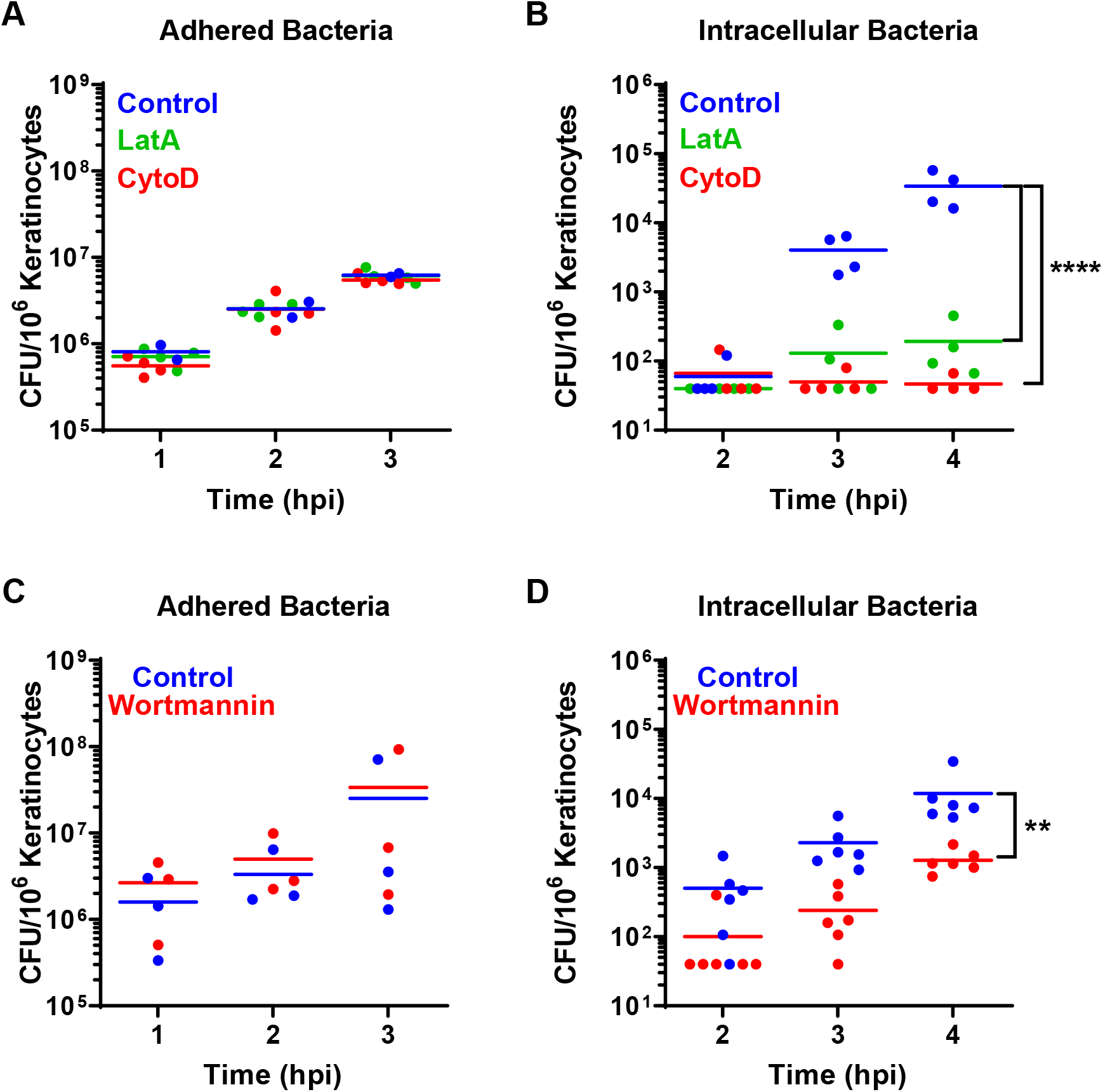
*E. faecalis* entry is dependent on actin polymerization. Keratinocytes were pre-treated with **(A**,**B)** actin inhibitors cytochalasin-D (CytoD 1 µg/ml) and latrunculin A (LatA 0.25 µg/ml) or **(C**,**D)** PI3K inhibitor wortmannin (0.1 mg/ml), followed by *E. faecalis* infection at MOI 100 for 1, 2, or 3 h. Host cells were lysed and associated adhered bacteria enumerated immediately after infection; or, to quantify intracellular CFU, the initial infection period was followed by 1 h antibiotic treatment, for a total of 2, 3 or 4 hpi, prior to lysis and enumeration. (**A**,**C**) Adherent or (**B**,**D**) intracellular bacteria were enumerated at the indicated time points (only significant differences are indicated). Solid lines indicate the mean for each data set of at least 3 independent experiments. **(B)** ****p<0.0001 2 way ANOVA, Tukey’s multiple comparisons test. **(D)** **p<0.01 2 way ANOVA, Sidak’s multiple comparisons test. (See **Supplementary Figure 3** for data related to the 24 hpi time point).

**Supplementary Figure 3 (related to Figure 3).**
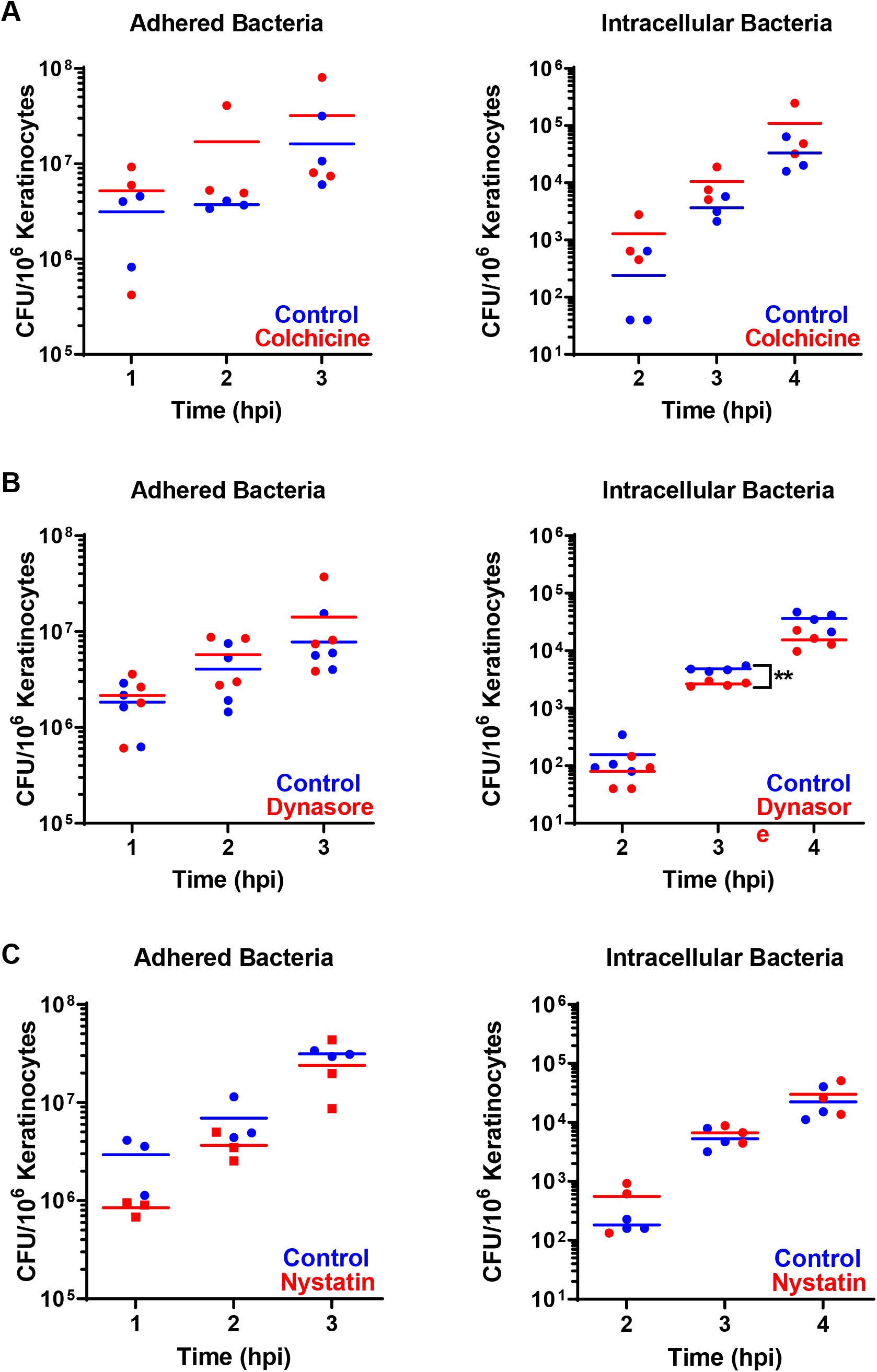
*E. faecalis* entry into keratinocytes is not dependent on microtubule polymerization, clathrin- and caveolae-mediated endocytosis. Keratinocytes were pre-treated with **(A)** microtubule inhibitor colchicine (10 µg/ml), **(B)** dynasore, an inhibitor of the large GTPase dynamin that is important for the formation of clathrin-coated vesicles (Kirchhausen, Macia, and Pelish 2008) (25 µg/ml), or **(C)** nystatin, which selectively affects caveolae-mediated endocytosis by binding sterols, causing caveolae and cholesterol disassembly in the plasma membrane (Ivanov 2008; Rothberg et al. 1992) (25 µg/ml). Treated cells were then infected with *E. faecalis* at MOI 100 for 1, 2, or 3 h. For enumeration of intracellular CFU, each infection period was followed by 1 h antibiotic treatment, for a total of 2, 3 or 4 hpi. (**A**,**C**) Adherent or (**B**,**D**) intracellular bacteria were enumerated at the indicated time points (only significant differences are indicated). Solid lines indicate the mean for each data set of at least 3 independent experiments. Note that we observed a slight decrease in the number of internalized bacteria across all time points in dynasore-treated cells, but we also found that dynasore kills >20% of all cells by 4 hpi (data not shown), suggesting that the reduction is associated with cell death and that dynamin is not essential for the uptake of *E. faecalis* into keratinocytes. **p<0.01 2 way ANOVA, Sidak’s multiple comparisons test.

### Intracellular *E. faecalis* traffics through early and late endosomes

Since we observed large numbers of *E. faecalis* within keratinocytes in the perinuclear region at 24 hpi, we hypothesized that *E. faecalis* may be trafficked through the host endo-lysosomal pathway. To interrogate this hypothesis, we infected cells for 4 hpi or 24 hpi and subsequently labeled the infected cells with fluorescently labelled antibody conjugates to visualize intracellular endo-lysosomal compartments: EEA1 (early endosome antigen 1, early endosome), LAMP1 (lysosomal-associated membrane protein, late endosome/pre-lysosome), M6PR (mannose-6-phosphate receptor, late endosome/pre-lysosome) and cathepsin D (lysosome) (**Figure 4** and **Supplementary Figure 4**). At 4 hpi, 64% (35/55) of internalized *E. faecalis* were found in LAMP1-positive and EEA1-negative late endosomes. Although EEA1-labeled early endosomes were often observed in close proximity to *E. faecalis*-containing compartments, only 5% (3/55) of internalized bacteria showed a clear association with EEA1, and these compartments did not contain LAMP1. The remainder (31% of internalized bacteria (17/55)) was neither positive for LAMP1 nor for EEA1 **(Figure 4A,B)**. These data suggest that internalized *E. faecalis* transits rapidly through early endosomes, reaching LAMP1-positive late endosomal compartments by 4 hpi.

**Figure 4.**
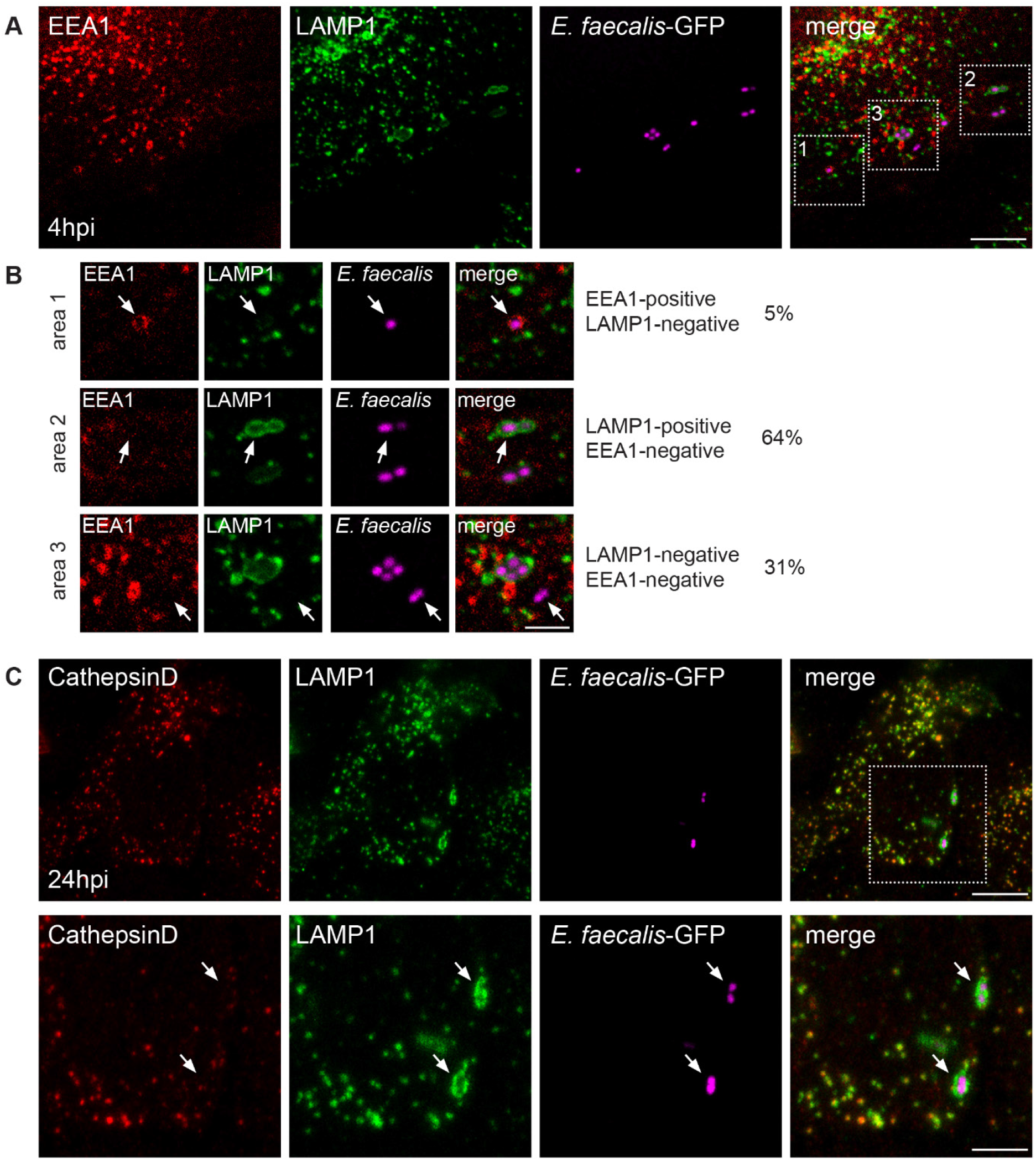
Internalized *E. faecalis* persist within a late endosomal compartment. **(A)** CLSM of infected HaCaTs stained with antibodies against EEA1 (early endosome) and LAMP1 (late endosome/lysosome) at 4 hpi. **(B)** Magnified images of boxed areas in (A) showing representative images of *E. faecalis* containing compartments. Measurements (percentages) are derived from 10 individual confocal images and a total of 55 *E. faecalis* diplococci. **(C)** CLSM of infected HaCaTs stained for the lysosomal protease cathepsin D (late endosome/lysosome) and LAMP1 (late endosome/lysosome) at 24 hpi. Bottom panel shows the boxed region above. Images are maximum intensity projections of 4-5 optical sections (∼2 µm z-volume) and are representative of 3 independent experiments. Scale bars: A: 10 µm; B: 2 µm; C top panel: 10 µm; C bottom panel: 5 µm. (See **Supplementary Figure 4** for additional images.)

While *E. faecalis* can survive in murine macrophages by resisting acidification, which in turn prevents fusion with lysosomes (Zou and Shankar 2016), this has not been previously documented in epithelial cells. Our results suggesting that *E. faecalis* can replicate in keratinocytes led us to hypothesize that fusion between late endosomes and lysosomes may be impeded, permitting intracellular survival. To determine if lysosomes fuse with *E. faecalis*–containing late endosomes, we co-stained infected cells at 24 hpi for LAMP1 and the lysosomal protease cathepsin D. We found that LAMP1 and cathepsin D colocalized in infected and non-infected cells, as expected. Interestingly, however, while 47% (30/64) of internalized *E. faecalis* were observed in LAMP1 positive compartments, these compartments were conspicuously devoid of cathepsin D (**Figure 4C**). The remainder (53% of internalized bacteria (34/64)) did not colocalise with either LAMP1 or cathepsin D. We also observed a complete lack of colocalization between *E. faecalis* and M6PR, which delivers lysosomal hydrolases to pre-lysosomal compartments (**Supplementary Figure 4**). Importantly, LAMP1 positive compartments containing *E. faecalis* often appeared distended, particularly at 24 hpi (**Figure 4C, Supplementary Figure 4**). Based on these observations, we propose that intracellular replication occurs within late endosomes until a bacterial threshold is reached, whereupon the compartment is unable to accommodate additional bacteria.

**Supplementary Figure 4 (related to Figure 4):**
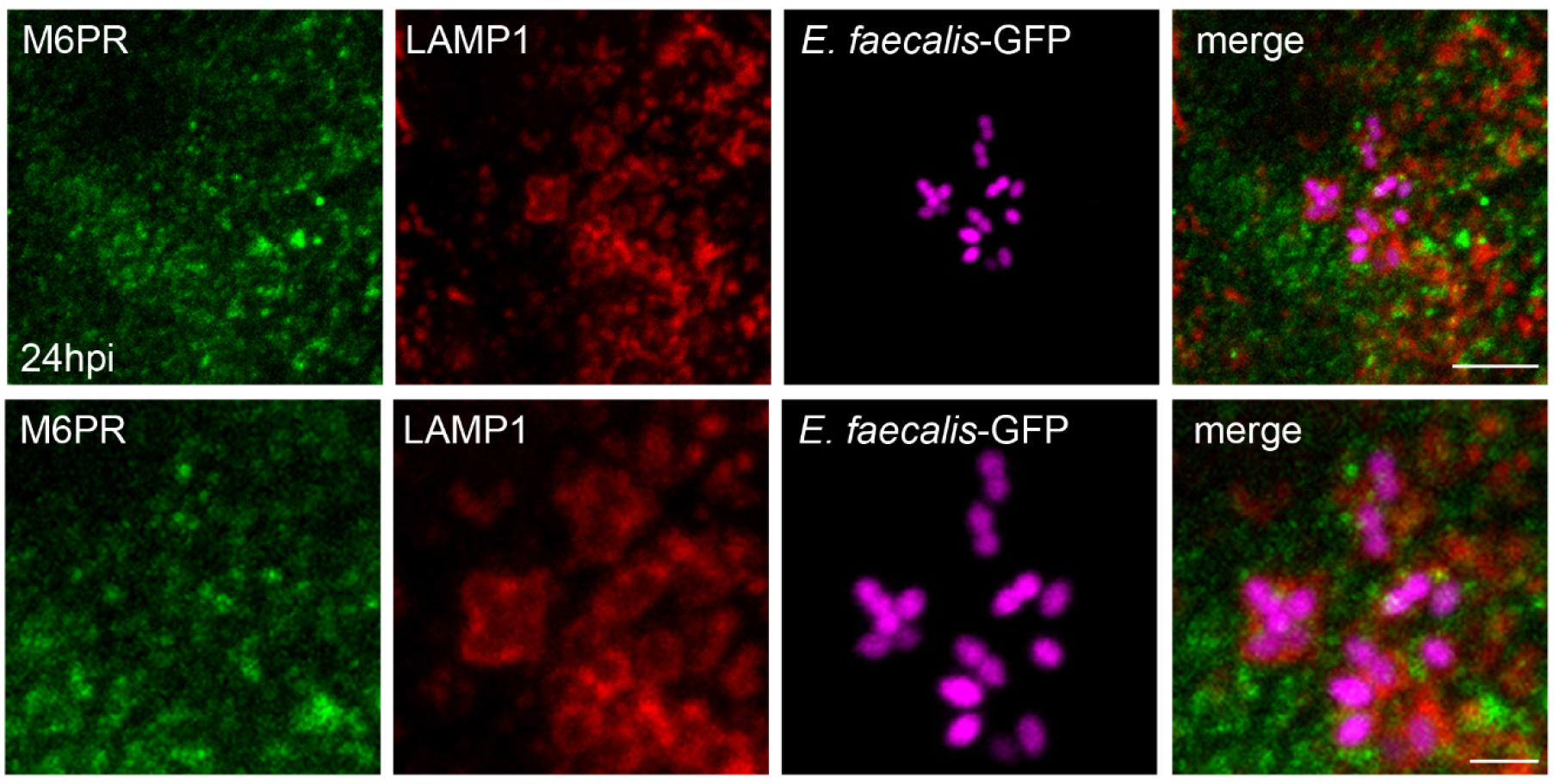
Internalized *E. faecalis* persist within late endosomal compartments. CLSM of infected HaCaTs stained with antibodies against M6PR (late endosome) and LAMP1 (late endosome/lysosome) at 24 hpi. Bottom panels show magnified images. Images are maximum intensity projections of 4-5 optical sections (∼2 µm z-volume) and are representative of 3 independent experiments. Scale bars are 5 µm in top panels and 2 µm in bottom panels.

### *E. faecalis* intracellular infection interferes with Rab5 and Rab7 protein levels

Rab5 and Rab7 are small GTPases that are critical for the formation of early and late endocytic compartments. Rab5 regulates early endosome/macropinosome formation while Rab7 is required for the maturation of early to late endosomes, as well as for the fusion of late endosomes with lysosomes (Wandinger-Ness and Zerial 2014). Late endosome-lysosome fusion is a critical step for the formation of an acidic and degradative compartment that eliminates bacteria, yet bacteria have evolved multiple mechanisms to interfere with this process (Mottola 2014). For instance, *Mycobacterium tuberculosis* affects Rab7 recruitment and, consequently, phagosome maturation, by interfering with Rab5 effectors (Saikolappan et al. 2012; Puri, Reddy, and Tyagi 2013). *Listeria monocytogenes* also affects Rab7 recruitment by inhibiting Rab5 GDP exchange activity in host cells (Prada-Delgado et al. 2005). Additionally, *Coxiella burnetii* can localize to Rab5 and LAMP1 positive compartments that lacks Rab7 (Ghigo et al. 2009; Ghigo, Colombo, and Heinzen 2012). To test whether *E. faecalis* infection affects Rab expression or localization in keratinocytes, we first performed CLSM on individual infected keratinocytes. We observed that Rab7 was associated with 28% (16/58) of *E. faecalis* containing compartments at 4 hpi and with 32% (10/31) of *E. faecalis* containing compartments at 24 hpi (**Figure 5A)**; however, some but not all LAMP1 positive *E. faecalis*-containing compartments were also Rab7 positive (**Supplementary Figure 6**), suggesting a degree of heterogeneity among the intracellular niche of *E. faecalis*. Next, we analyzed the expression levels of Rab7 and Rab5 in the infected keratinocyte population, following infection with either *E. faecalis* strain OG1RF or strain V583 (**Figure 5B,C**). Somewhat unexpectedly, although we observed Rab7 associated with some *E. faecalis* compartments, we observed a global reduction in both Rab7 and Rab5 protein levels for both strains at 4 hpi. While Rab7 levels were restored by 24 hpi for both strains, Rab5 in V583-infected keratinocytes remained low at 24 h, which may correlate with greater CFU and intracellular survival for V583 compared to OG1RF (**Supplementary Figure 1C,D**). The expression of other endo-lysosomal proteins, such as cathepsin D, were unchanged upon *E. faecalis* infection (**Supplementary Figure 6**), indicating that *E. faecalis* selectively interferes with the functions of endosomal Rab proteins. Taken together, the combined *E. faecalis*-mediated reduction in Rab expression, coupled with the ability of nearly 70% of *E. faecalis*-containing compartments to avoid Rab7 recruitment could explain the lack of colocalization of cathepsin D with *E. faecalis*-containing compartments (**Figure 4C**) and suggests that these compartments do not fuse with lysosomes.

**Figure 5.**
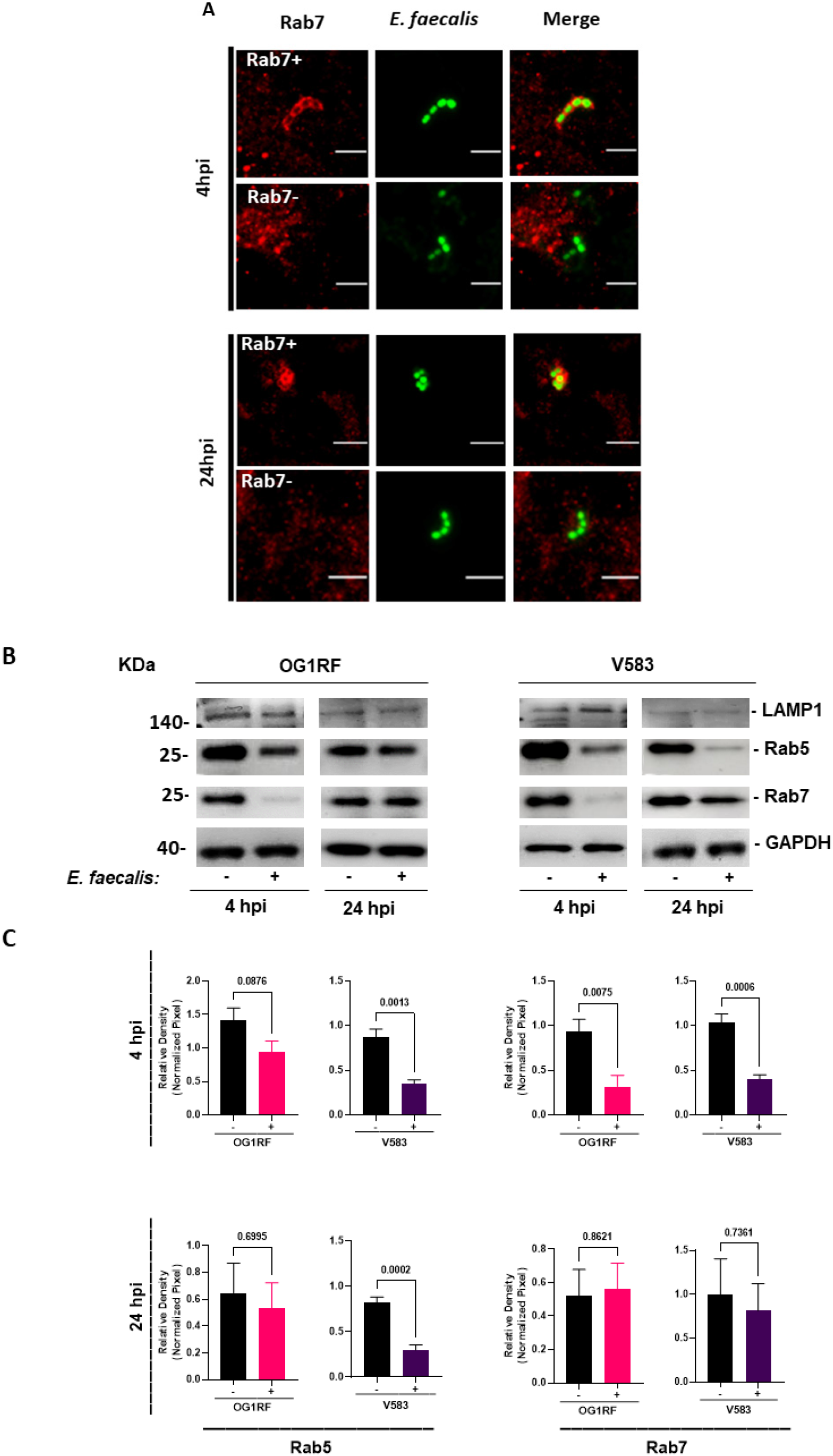
Internalized *E. faecalis* reduces expression of both Rab5 and Rab7 at 4 hpi. **(A)** CLSM of infected HaCaTs with fluorescent labelling of Rab7 (late endosome) and fluorescent *E. faecalis* (Dasher). Images show examples of Rab7+ and Rab7-compartments. Images shown are representative of 3 independent experiments. Scale bar: 5 µm. **(B)** Whole cell lysate analyzed by immunoblotting with antibodies α-LAMP1, α-Rab5, α-Rab7 and α-GAPDH. HaCaT cells were incubated with (+) and without (-) *E. faecalis* OG1RF and V583 for 4 h and 24 h. **(C)** Relative density of the bands of interest were normalized against loading control (GAPDH). Error bars represent biological replicates and mean ± SEM from at least 3 independent experiments. Statistical analysis was performed using unpaired T-test with Welch’s correction.

**Supplementary Figure 5 (related to Figure 5).**
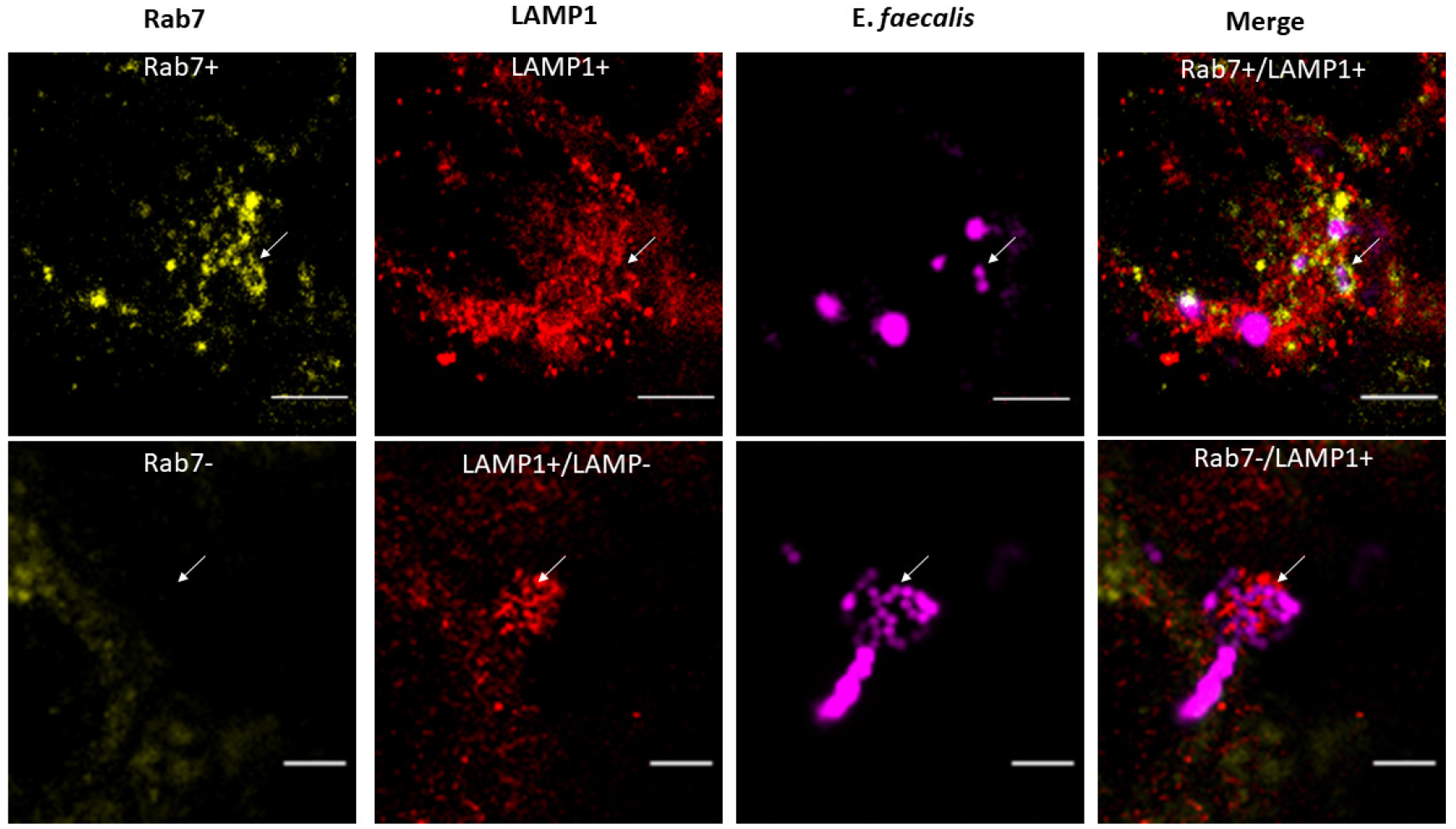
*E. faecalis* is found in heterogeneously labelled Rab7/LAMP1 compartments. CLSM of infected HaCaTs with fluorescent labelling of Rab7 and LAMP1 (late endosome) and fluorescent *E. faecalis* (Dasher). Images show examples of Rab7+/LAMP1+ (Top panel), LAMP1+/Rab7- and LAMP1+/Rab7- (Bottom panel) compartments. Images shown are representative of 3 independent experiments. Scale bar: 5 µm.

**Supplementary Figure 6 (related to Figure 5).**
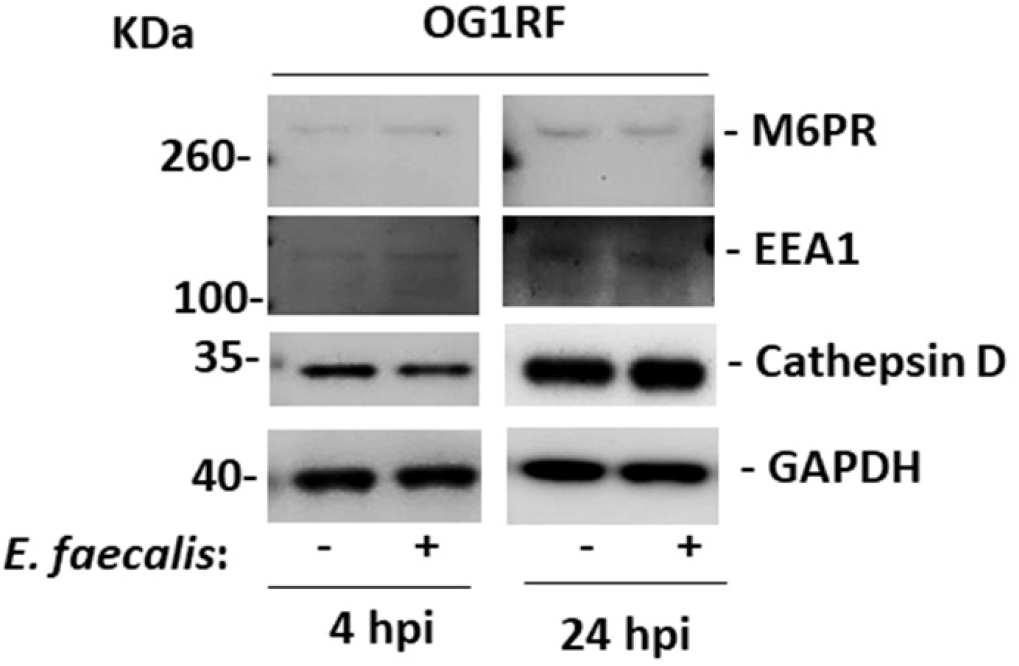
*E. faecalis* infection of keratinocytes does not alter expression of other endosomal proteins. Whole cell lysates analyzed by immunoblot with antibodies α-M6PR, α-EEA1, α-cathepsinD, and α-GAPDH. HaCat cells were incubated with (+) and without (-) *E. faecalis* OG1RF for 4 h and 24 h. Images shown are representative of 3 biological replicates.

### *E. faecalis* containing vacuoles do not fuse with lysosomes

To visualize the association between *E. faecalis*-containing compartments and endo-lysosomal organelles with greater resolution, we turned to correlative light and electron microscopy (CLEM). We created a HaCaT cell line that stably expresses LAMP1-mCherry and infected these cells with GFP-expressing *E. faecalis* for 18 hours (3 hours followed by 15 hours antibiotic treatment). Confocal microscopy of fixed cells enabled us to locate *E. faecalis* and LAMP1+ compartments in infected cells before processing them for serial section transmission electron microscopy (TEM) (**Figure 6, Supplementary Figure 7**). These experiments revealed several features of *E. faecalis* intracellular infection that we could not appreciate using fluorescence microscopy alone. First, we observed *E. faecalis* in LAMP1 positive compartments as well as in vacuoles that appeared to be devoid of LAMP1 **(Figure 6A-C**), which is in line with our immuno-fluorescence microscopy data **(Figure 4, Supplementary Figure 4)**. Second, and importantly, regardless of the degree of colocalisation with LAMP1, *E. faecalis-*containing vacuoles were invariably bounded by a single membrane (**Figure 6D-H**). In addition, most internalized bacteria appeared to be morphologically intact, with a uniform density and a clearly defined septum and bacterial envelope (**Figure 6D-H**). Third, we did not find evidence of multiple replicating *E. faecalis* within a single LAMP1 positive compartment leading to membrane distension, as predicted from immuno-fluorescence imaging (**Figure 4B, Supplementary Figure 4**). Rather, we observed at most two diplococci within a single compartment (**Figure 6D**). Finally, although lysosomes (defined as LAMP1 positive organelles with a dense ultrastructural appearance) were often located in close proximity to *E. faecalis* containing vacuoles, we did not observe any obvious fusion events between the two compartments (**Figure 6C, 5E**). In some instances, however, vacuoles harbouring *E. faecalis* appeared to contain LAMP1 positive multi-lamellar bodies (MLBs) **(Figure 6G, Supplementary Figure 7)**. Together, these EM data confirm both that internalized *E. faecalis* can survive in late endosomal organelles and that there is heterogeneity within the intracellular niches for this organism. Further, *E. faecalis-*containing vacuoles do not exhibit lysosomal features and do not appear to fuse with lysosomes. These findings raise the possibility that *E. faecalis* could be hijacking the endo-lysosomal pathway, altering organelle identity to prevent lysosomal recognition, and allowing for intracellular survival, replication and eventual escape.

**Figure 6:**
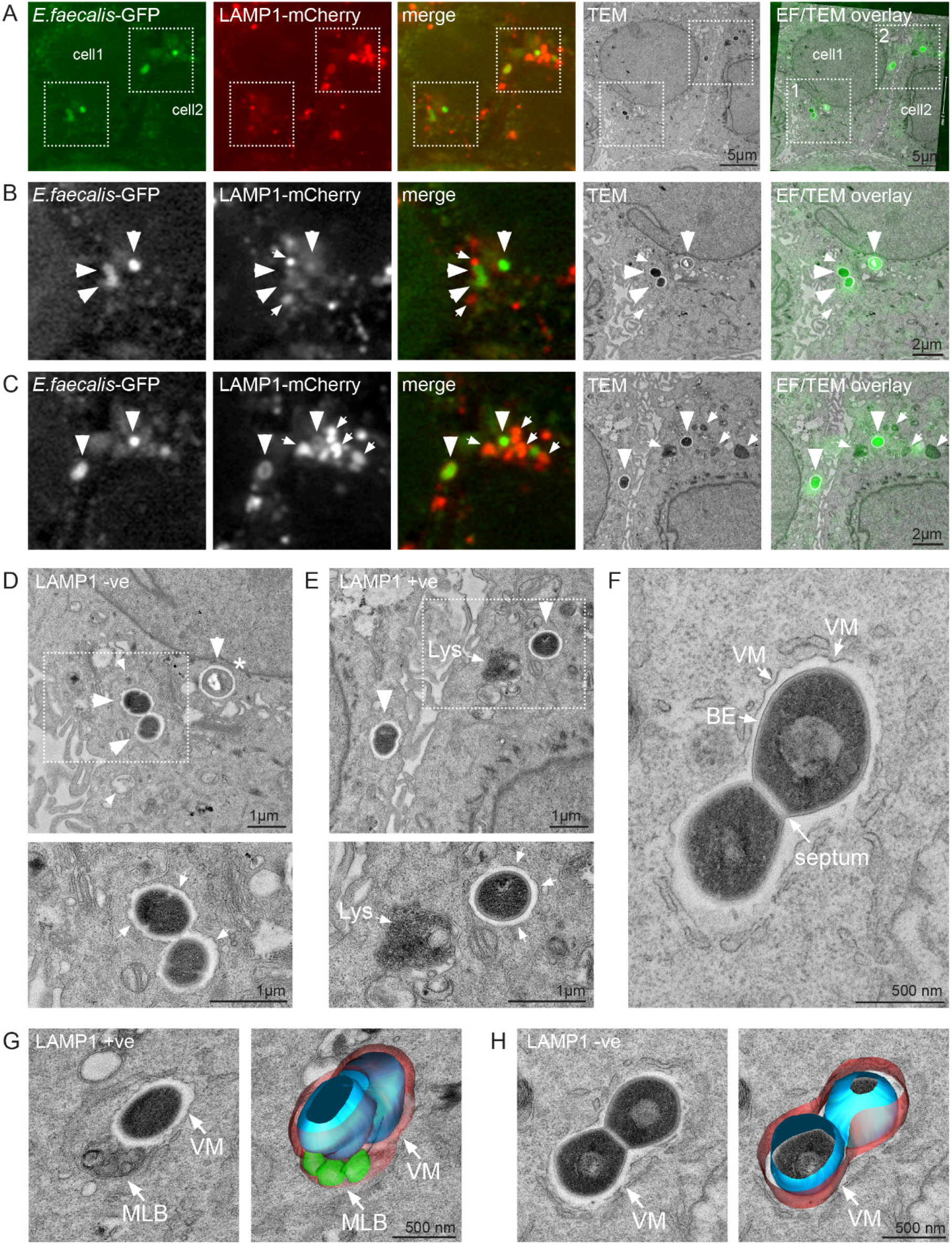
Correlative light and electron microscopy of *E. faecalis* infected keratinocytes. **(A)** Spinning disk confocal microscopy and correlative TEM of HaCaTs stably expressing LAMP1-mCherry infected with *E. faecalis-*GFP at 18 hpi. Confocal images are maximum intensity projections of 4-5 optical sections (∼2 µm z-volume). **(B and C)** Enlarged views of the two areas highlighted in (A); panels B and C show the boxed areas 1 and 2 in (A), respectively. Large arrowheads indicate *E. faecalis* containing vacuoles, small arrows indicate LAMP1-positive (LAMP1+ve) compartments. Note that *E. faecalis* is present in LAMP1 negative (LAMP1-ve) vacuoles in (B), and in LAMP1+ve vacuoles in (C). Note that LAMP1+ve compartments in (C) exhibit ultrastructural features typical of lysosomes. * indicates a bacterium with altered appearance. **(D and E)** Representative high magnification TEM images of LAMP1-ve (D) and LAMP1+ve (E) *E. faecalis* containing vacuoles. Images in (D) and (E) correspond to data shown in (B) and (C), respectively. A LAMP1+ve lysosome (Lys) in close proximity to an *E. faecalis* containing vacuole is highlighted in (E). Arrowheads in the lower panels indicate the presence of a single layer membrane surrounding the bacterial vacuole. **(F)** High magnification view of an *E. faecalis* containing vacuole. The vacuolar membrane (VM), the bacterial envelope (BE), and the septum are indicated. **(F and G)** 3D surface rendering of representative *E. faecalis* containing vacuoles reconstructed from serial TEM sections. An *E. faecalis* containing vacuole containing a LAMP1+ve multilamellar body (MLB) is shown in (G), while the vacuole shown in (F) is LAMP1-ve and does not contain a MLB. (See **Supplementary Figure 7** for data related to F-H).

**Supplementary Figure 7 (related to Figure 6):**
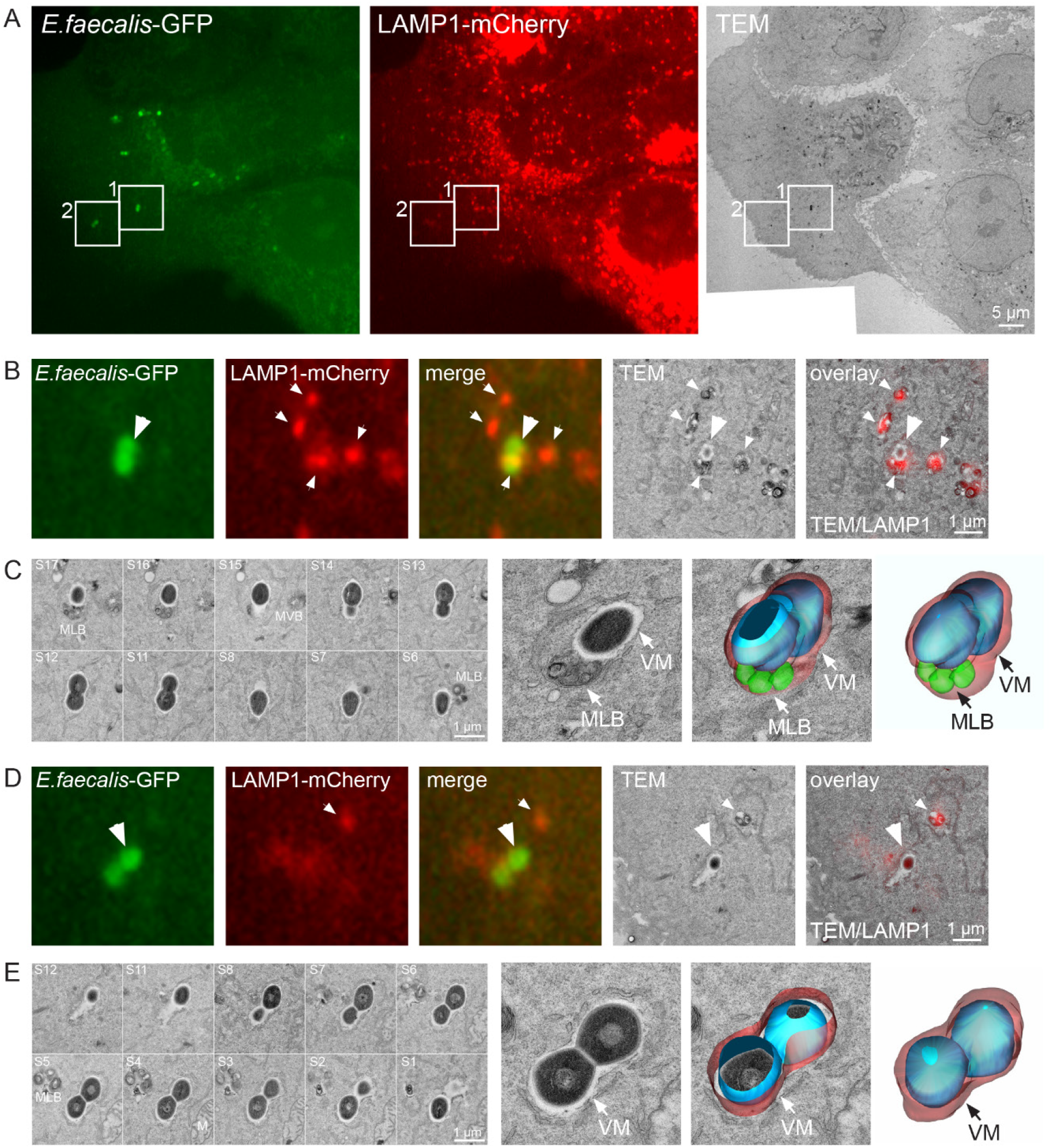
Correlative light and electron microscopy of *E. faecalis* infected keratinocytes. **(A)** Spinning disk confocal microscopy and correlative TEM of HaCaTs stably expressing LAMP1-mCherry infected with *E. faecalis-*GFP at 18 hpi. Confocal images are maximum intensity projections of 4-5 optical sections (∼2 µm z-volume). (B) Enlarged views of area 1 highlighted in (A). (C) Serial section TEM and 3D surface rendering of the area shown in (B). (D) Enlarged views of area 2 highlighted in (A). (E) Serial section TEM and 3D surface rendering of the area shown in (D). Large arrowheads indicate *E. faecalis* containing vacuoles, small arrows indicate LAMP1-positive compartments. VM: vacuolar membrane (VM); MLB: multilamellar body. MLB: multilamellar body. An *E. faecalis* containing vacuole containing a LAMP1+ve MLB is shown in (B and C), while the *E. faecalis* containing vacuole shown in (D and E) is LAMP1-ve and does not contain a MLB (data pertinent to Figure 5F-H).

### Intracellular *E. faecalis* is primed for more efficient reinfection

To investigate whether internalization of *E. faecalis* into keratinocytes provides an advantage for subsequent reinfection, we harvested intracellular bacteria and measured its ability to reinfect keratinocytes. An initial infection was performed at MOI 50 for 3 hours to isolate intracellular bacteria. Intracellular-derived bacteria were then used for reinfection of keratinocytes at an MOI of 0.1, the highest MOI practically attainable given the low intracellular CFU, for another 3 hours. 1 hour of gentamicin/penicillin treatment was performed after both the initial infection and the second round of infection. Internalization recovery ratios were determined by comparing inoculum CFU to intracellular CFU bacteria during the reinfection assay. Parallel experiments with *E. faecalis* not yet exposed to keratinocytes at comparable MOI showed that reinfection with intracellular-derived bacteria resulted in significantly higher internalization rates, as shown by the recovery ratio **(Figure 7A)**. Intracellular growth exclusively promoted reinfection, because total cell associated bacteria comprising both adherent and intracellular bacteria, was not significantly different from a planktonically grown inoculum **Figure 7B)**. These results are similar to observations made in *S. pyogenes*, where longer periods of internalization in macrophages increased recovered CFU during subsequent reinfections (Hertzen et al. 2012). Taken together, these data suggest that internalized *E. faecalis* can more efficiently reinfect host cells.

**Figure 7.**
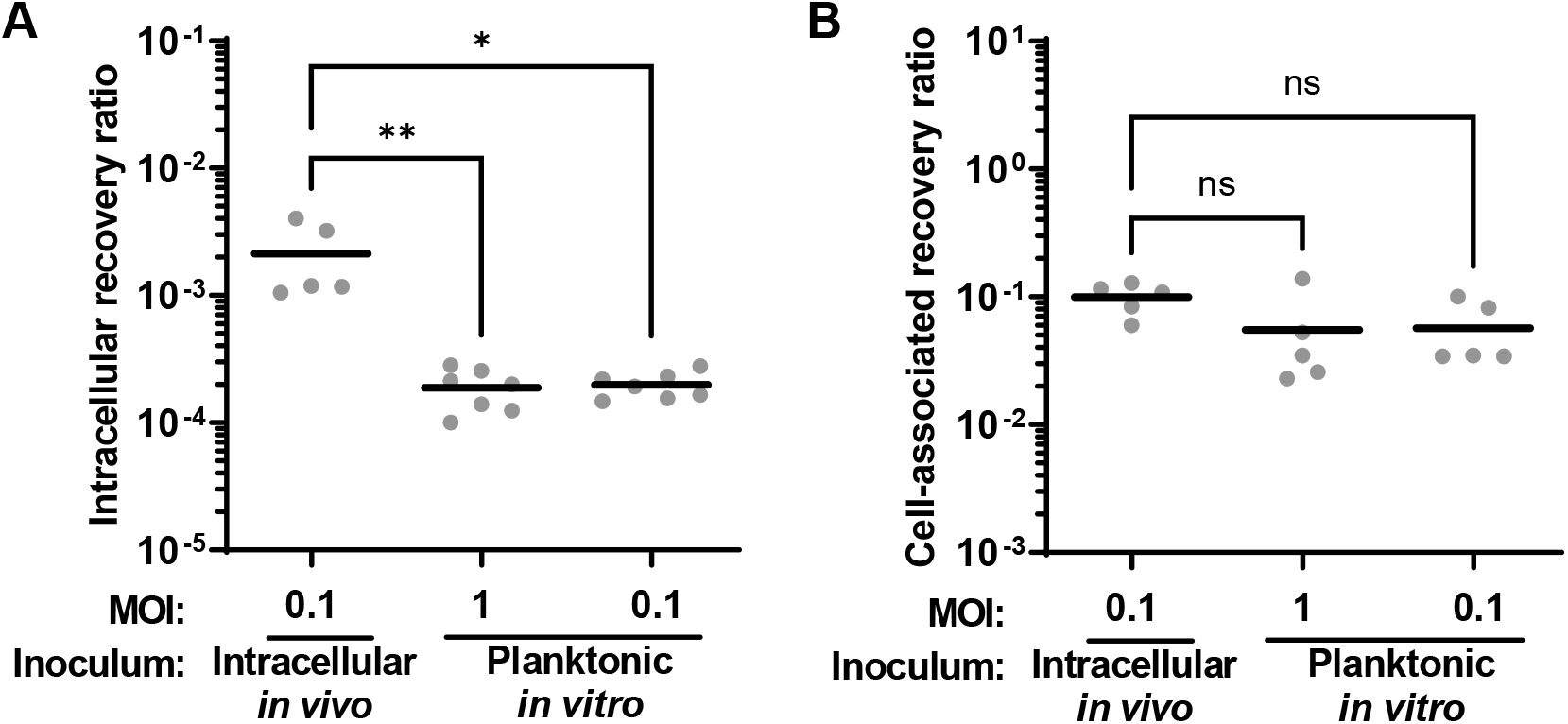
Increased recovery of internalized *E. faecalis* upon reinfection of keratinocytes by intracellular bacteria. Intracellular bacteria were isolated from infected keratinocytes and used as the inoculum (intracellular *in vivo*) for reinfection of new monolayers of keratinocytes. Parallel infections at various MOI were performed using *E. faecalis* not yet exposed to keratinocytes, grown planktonically *in vitro* as the inoculum. **(A)** Infections proceeded for 3 h followed by 1 h of antibiotic exposure to kill the extracellular bacteria prior to intracellular CFU enumeration. **(B)** Infections proceeded for 3 h prior extensive washing to remove non-adhered extracellular bacteria, prior to enumeration of total cell-associated (both extracellular adhered and intracellular) CFU. Each circle represents CFU data averaged from 3 separate wells from a single biological experiment, showing a total of 5-7 independent experiments. Data are represented in the figure as the recovery ratio, which for **(A)** intracellular is the CFU recovered divided by the inoculum CFU and for **(B)** cell-associated is the CFU recovered divided by the non-cell-associated extracellular CFU in the same well, to account for cell growth during the assay. Horizontal black line indicates the mean for each condition. *p<0.05, **p<0.01 Kruskal Wallis test with Dunn’s post test.

## Discussion

*E. faecalis* is among the commonly isolated microbial species cultured from chronic wound infections. The ability of *E. faecalis* to persist in the face of a robust immune response and antibiotic therapy is frequently attributed to its ability to form biofilms during these infections. However, a number of bacterial pathogens undertake an intracellular pathway during infection that can contribute to persistent and or recurrent infection. This is well-described for uropathogenic *E. coli*, which can replicate to high numbers within urothelial cells as intracellular bacterial communities, or can persist in a quiescent intracellular state within LAMP1 positive compartments for long periods of time, promoting recurrent and chronic infection (Mysorekar and Hultgren 2006; Klein and Hultgren 2020; Anderson et al. 2003; Guiton et al. 2012). While there are numerous reports of intracellular *E. faecalis* within a variety of non-immune cells (Bertuccini et al. 2002; Horsley et al. 2018; Horsley et al. 2013; Olmsted et al. 1994; Wells, Jechorek, and Erlandsen 1990; Wells et al. 1988; Baldassarri et al. 2005; Millan et al. 2013), the contribution of an intracellular lifecycle to *E. faecalis* infection has been minimally investigated. Here, we report that *E. faecalis* become internalized into keratinocytes via macropinocytosis, whereupon they manipulate the endocytic-lysosomal pathway, enabling their replication and survival. These findings raise the possibility that this intracellular lifecycle may be linked to persistent and chronic infections, such as those that occur in wounds. Further, we demonstrate that intracellularity may be physiologically relevant in a mouse model of wound infection, where *E. faecalis* exists within both immune and non-immune cells for at least 5 days after infection. Importantly, *E. faecalis* recovered from within keratinocytes are primed to more efficiently infect new keratinocytes to seed another round of infection.

Previous studies using either professional or non-professional phagocytic cell lines have reported the internalization, but not the replication of intracellular *E. faecalis* (Gentry-Weeks et al. 1999; Bertuccini et al. 2002; Baldassarri et al. 2005) as these studies used only antibiotic protection assays coupled with TEM at single time points. Here, we performed antibiotic protection assays coupled with imaging across multiple time points, and our results similarly show that *E. faecalis* can enter and survive intracellularly up to 72 hpi. Importantly, imaging data in this study obtained at 4 hpi compared to 24 hpi showed that infected cells contain more bacteria at later time points, supporting the conclusion that *E. faecalis* can multiply within keratinocytes. This is the first reported evidence, to our knowledge, of *E. faecalis* intracellular replication within epithelial cells. Consistent with our observation, *E. faecalis* has also been shown to replicate within human hepatocytes *in vitro* and has been observed as clusters in association with hepatocytes in a mouse model of intravenous infection (Pascale Serror, personal communication). Together these data suggest that once *E. faecalis* enters host epithelial cells, at least some of the bacteria are able to replicate within those cells. Other “classical” extracellular bacteria including *S. aureus* and *P. aeruginosa* are also able to replicate intracellularly (Flannagan, Heit, and Heinrichs 2016; Jolly et al. 2015; Garzoni and Kelley 2009; Jarry and Cheung 2006; Qazi et al. 2004; Mukherjee, Khatua, and Mandal 2020). Similar studies have also shown that *S. pyogenes* can be taken up by both immune and non-immune cells, where it can replicate, survive host defenses and disseminate to distant sites (Bastiat-Sempe et al. 2014; Osterlund and Engstrand 1997). Given that non-professional phagocytes often lack the ability to produce reactive oxygen species (ROS) or cytokines in response to infection (Rabinovitch 1995), it is therefore possible that keratinocytes are less able to eradicate internalized bacteria and may provide a safe haven for *E. faecalis*.

In this work, we show that *E. faecalis* enters keratinocytes in a process that is dependent on actin polymerization and PI3K signalling, and independent of receptor (clathrin)- or caveolae-mediated endocytosis. Chemical inhibition of actin polymerization by cytochalasin D and PI3K signaling by wortmannin specifically affects macropinocytosis but not receptor (clathrin)-mediated endocytosis (Araki, Johnson, and Swanson 1996; Gaidarov et al. 1999; Swanson and Watts 1995). These findings suggest that *E. faecalis* strain OG1RF enters keratinocytes in a macropinocytotic process. A previous study suggested that clinical isolates of *E. faecalis* enter into HeLa (human epithelioid carcinoma) cells via either macropinocytosis or clathrin-mediated endocytosis, supported by inhibitors of microtubule polymerization and cytosolic acidification that reduced intracellular CFU (Bertuccini et al. 2002). It may be that different strains of *E. faecalis* enter mammalian cells by different mechanisms, and *E. faecalis* OG1RF used in this study preferentially enters via macropinocytosis. However, another study reported that *E. faecalis* OG1 strain derivatives, closely related to OG1RF, entered human umbilical vein endothelial cells (HUVEC) cells via receptor (clathrin)-mediated endocytosis, in a cytocholasin D- and colchicine-dependent manner (Millan et al. 2013). Because Millan et al used similar drug concentrations as we did, we suggest that OG1-related strains may enter epithelial cells via macropinocytosis and endothelial cells via receptor (clathrin)-mediated endocytosis.

Additionally, once inside keratinocytes, *E. faecalis* commences trafficking through the endosomal pathway. At 4 hpi, most internalized *E. faecalis* were observed in the periphery of infected cells. At this time point, a small percentage of internalized *E. faecalis* colocalized with the early endosomal marker EEA1, while the majority were in compartments that were heterogeneously positive for the late endosomal markers LAMP1 and/or Rab7. These data suggest that internalized *E. faecalis* is taken up into EEA1-positive early endosomes/macropinosomes, yet rapidly transits into late endosomal compartments. Interestingly, at 4 hpi, Rab5 and Rab7 protein levels in infected keratinocytes were markedly decreased in comparison to non-infected keratinocytes. We predict that *E. faecalis* infection-driven reduction in Rab expression is crucial to determine the outcome of *E. faecalis* intracellular survival since Rab5 and Rab7 control important fusion events between early and late endosomes and late endosomes and lysosomes (Mottola 2014). Rab GTPases are commonly hijacked by bacteria to promote their survival (Spano and Galan 2018). For comparison, intracellular microbes such as *M. tuberculosis* and *L. monocytogenes* distinctively modify the Rab5 machinery arresting phagosome maturation (Mottola 2014). *C. burnetii* prevents Rab7 recruitment (Barry et al. 2012) and *B. cenocepacia* affects Rab7 activation (Huynh et al. 2010). However, to the best of our knowledge, our data are the first to show a decrease in overall Rab5 and Rab7 protein levels as a potential bacterial subversion mechanism for the macropinosome. Studies are underway to determine the bacterial factors and mechanisms by which *E. faecalis* affects Rab protein levels. By 24 hpi, most *E. faecalis* in infected cells were in the perinuclear region. While *E. faecalis*-containing LAMP1-positive compartments appeared distended at 24 hpi, many *E. faecalis* were not tightly associated with LAMP1 or Rab7. Furthermore, we observed no cathepsin D in any *E. faecalis*-containing compartment, suggesting that late endosomes containing internalized bacteria could be missing markers or that these late endosomal compartments have been modified, making lysosomal fusion impossible. Other intracellular pathogens such as *C. burnetii* and *Francisella tularensis* also reside in compartments devoid of cathepsin D, or in compartments with very low levels of cathepsin D (Ghigo et al. 2002; Clemens, Lee, and Horwitz 2004). The authors suggest that this was achieved by escaping fusion with lysosomes. In support of this view, TEM revealed that all membrane-bound *E. faecalis* were spatially separated from lysosomes, and there was no indication of membrane fusion between *E. faecalis-*containing compartments and lysosomes. Notably, we observed some *E. faecalis* in association with LAMP1 and Rab7, suggesting that some internalized *E. faecalis* cells may transit via the normal endocytic pathway and fuse with lysosomes. Collectively our results support a model in which late endosomes containing *E. faecalis* are modified, preventing the expected destruction of intracellular *E. faecalis* by lysosomal fusion and allowing them to replicate from within.

We propose three potential fates for internalized *E. faecalis*. 1) Macropinosome maturation into Rab7/LAMP1-positive late endosomes and fusion with the lysosome, leading to the degradation of *E. faecalis* in a small subset of infected cells (**Figure 8-I**). 2) Macropinosome maturation into LAMP1-positive but Rab7-negative compartments, leading to *E. faecalis* survival (**Figure 8-II**). 3) Macropinosome maturation into compartments lacking both Rab7 and LAMP1, which would also lead to *E. faecalis* survival (**Figure 8-III**). Furthermore, we cannot exclude the possibility that *E. faecalis*-containing compartments initially contain both late endosomal markers, but are subsequently modified by *E. faecalis* to increase its survival. In other words, there could be a transition from **I** into **II** and/or **III** mediated by downregulation of Rab5 and Rab7. Finally, we have also shown that infected host cells can eventually die, releasing *E. faecalis* into the periphery of dead host cells. Based on reinfection studies, we propose that those bacteria released from dead cells may be primed to infect other cells, resulting in an enhanced cycle of reinfection. Altogether, our work has demonstrated that *E. faecalis* can enter, survive, replicate and escape from keratinocytes *in vitro*. If this intracellular lifecycle also exists *in vivo*, and extends to other cell types such as macrophages as our data suggest, these findings may allow for an abundant protective niche for bacterial persistence that could contribute to the chronic persistent infections associated with *E. faecalis*.

**Figure 8.**
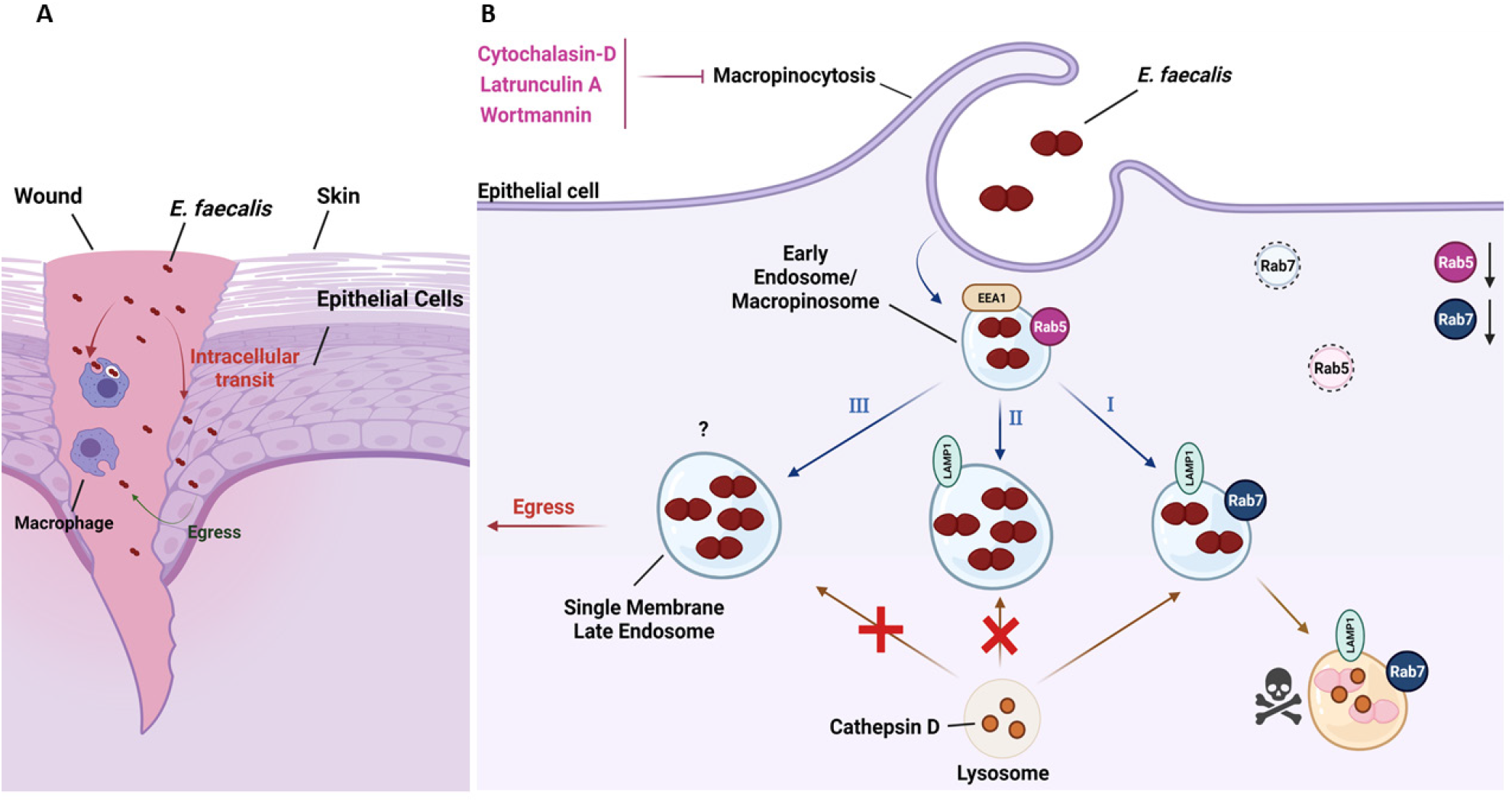
The intracellular lifestyle of *E. faecalis*. **(A)** *E. faecalis* can survive intracellularly either inside keratinocytes or macrophages and contribute to reinfection in the wound site. **(B)** *E. faecalis* is taken up by keratinocytes via macropinocytosis. Treatment of keratinocytes with macropinocytosis inhibitors such as cytochalasin D, latrunculin A and wortmannin prevents *E. faecalis* uptake. **(I)** *E. faecalis* was observed inside single membrane endosomes that were positive to different endosomal proteins indicating that *E. faecalis* may transit through the normal endocytic pathway inside keratinocytes. **(II and III)** *E. faecalis* interferes with Rab5 and Rab7 proteins levels which could help prevent the late endosome to fuse with lysosome. The low percentage of compartments positive to Rab7 combined with the varied percentages of LAMP1+ compartments indicates that *E. faecalis* may be affecting the expected macropinosome transit contributing to *E. faecalis* intracellular survival. Intact Rab5 and Rab7 proteins are shown in full color while affected ones are depicted as degraded in light color. Created with BioRender.com.

## Experimental Procedures

### Bacterial strains and growth conditions

*Enterococcus faecalis* strain OG1RF (ATCC47077) and its derivative strains, including OG1RF strain SD234 which chromosomally expresses GFP (Debroy et al. 2012) or fluorescent *E. faecalis* (Dasher) (built using the IP-Free©Fluorescent ProteinPaintbox™ -*E*. *coli* (Hallinen, Wittman, and Aukema 2020)), were grown using Brain Heart Infusion (BHI) (Becton, Dickinson and Company, Franklin Lakes, NJ). Unless otherwise stated, bacterial strains were streaked from glycerol stocks stored at - 80°C, inoculated and grown overnight statically for 16-20 h in 20 ml of liquid BHI broth. Cells were harvested by centrifugation at 5000 RPM (4°C) for 4 min. The supernatant was discarded, and the pellet washed with 1 ml of sterile phosphate buffered saline (PBS). The pellet was then resuspended in sterile PBS to an optical density (OD_600nm_) of 0.7 for *E. faecalis*, equivalent to 2–3×10^8^ colony forming units (CFU).

### Mouse wound excisional model

Mouse wound infections were modified from a previous study (Chong et al. 2017). Briefly, male wild-type C57BL/6 mice (7-8 weeks old, 22 to 25 g; InVivos, Singapore) were anesthetized with 3% isoflurane. Following dorsal hair trimming, the skin was then disinfected with 70% ethanol before creating a 6-mm full-thickness wound using a biopsy punch (Integra Miltex, New York, USA). *E. faecalis* corresponding to 2-3 × 10^6^ CFU was added to the wound site and sealed with a transparent dressing (Tegaderm™ 3M, St Paul Minnesota, USA). At the indicated time points, mice were euthanized and a 1 cm by 1 cm squared piece of skin surrounding the wound site was excised and collected in sterile PBS. Skin samples were homogenized, and the viable bacteria enumerated by plating onto both BHI plates and rifampicin (50 ug/ml) selection plates to ensure all recovered CFU correspond to the inoculating strain.

### Fluorescence-activated cell sorting (FACS)

Excised skin samples were harvested, placed in 1.5 ml Eppendorf tubes containing 2.5 U/ml liberase prepared in DMEM (10% FBS) with 500 μg/mL of gentamicin/penicillin G (Sigma-Aldrich, St. Louis, MO) and minced with surgical scissors. The mixture was then transferred into 6-well plates and incubated for 1 h at 37°C in a 5% CO2 humidified atmosphere with constant agitation. Dissociated cells were then passed through a 70 μm cell strainer to remove undigested tissues and spun down at 1350 RPM for 5 min at 4°C. The enzymatic solution was then aspirated, and cells were blocked in 500 μl of FACS buffer (2% FBS (Gibco, Thermo Fisher Scientific, Singapore), 0.2mM ethylenediaminetetraacetic acid (EDTA) in PBS (Gibco, Thermo Fisher Scientific, Singapore). Cells were then incubated with 10μl of Fc-blocker (anti-CD16/CD32 antibody) for 20 min, followed by incubation with an anti-mouse CD45-Cy7 conjugated antibody (BD Pharmingen™, Singapore) (1:400 dilution) for 20 min at room temperature. Cells were then centrifuged at 1350 RPM for 5 min at 4°C and washed in FACS buffer before a final resuspension in FACS buffer. Following which, cells were then sorted using a BD FACSAria™ 3 sorter, equipped with 4 air-cooled lasers (355 nm UV, 488 nm Blue, 561 nm Yellow/Green and 633 nm Red) (Becton Dickinson, Franklin Lakes, NJ). Post-sorting, cells were lysed with 0.1% Triton–X100 (Sigma-Aldrich, St. Louis, MO) and intracellular bacteria plated onto both BHI plates and antibiotic selection plates to ensure all recovered CFU correspond to the inoculating strain. Supernatants were also plated to ensure no bacteria was present post-sorting.

### Cell culture

The spontaneously immortalized human keratinocyte cell line, HaCaT (AddexBio, San Diego, CA) was cultured at 37°C in a 5% CO2 humidified atmosphere. Cells were grown and maintained in Dulbecco’s modified Eagle’s medium (DMEM) (Gibco; Thermo Fisher Scientific, Singapore) with 10% heat-inactivated fetal bovine serum (FBS) (PAA, GE Healthcare, Singapore), and 100 U of penicillin–streptomycin (Gibco, Thermo Fisher Scientific, Singapore) where appropriate for extracellular bacterial killing. The culture medium was replaced once every three days, and upon reaching 80% confluency, cultures were passaged. Passaging was achieved by treatment with 0.25% trypsin-EDTA (Gibco; Thermo Fisher Scientific, Singapore) for 6 min and seeding cells at a density of 2×10^6^ cells/T75 flask (Nunc; Thermo Fisher Scientific, Singapore).

### Intracellular infection assay

Keratinocytes were seeded at a density of 5×10^5^ cells/well in a 6-well tissue culture plate (Nunc; Thermo Fisher Scientific, Singapore) and grown for 3 days at 37°C in a 5% CO2 humidified atmosphere. After 3 days, each well had approximately 1– 1.5×10^6^ keratinocytes. Keratinocytes were infected at a multiplicity of infection (MOI) of 100, 10 or 1 for up to 3 h. Following infection, the media was aspirated, and the cells washed three times in PBS and either lysed in 0.1% Triton–X100 (Sigma-Aldrich, St. Louis, MO) for enumeration of cell-associated/adhered bacteria, or incubated with 500 μg/ml of gentamicin/penicillin G (Sigma-Aldrich, St. Louis, MO) in complete DMEM for 1-70 h to selectively kill extracellular bacteria. The antibiotic containing medium was then removed, the cells washed 3 times in PBS, the intracellular bacteria enumerated.

### Chemical inhibition of endocytosis

All chemical inhibitors were purchased from Sigma-Aldrich (St. Louis, MO), unless otherwise stated. Stock solutions of cytochalasin D (1 mg/ml), latrunculin A (100 μg/ml), colchicine (10 mg/ml), dynasore (25 mg/ml), nystatin (25 mg/ml) and wortmannin (10 mg/ml) were dissolved in DMSO unless otherwise indicated and stored at − 20°C. Pharmacological inhibitors were added to cells 30 min prior to any infection and maintained throughout the course of the infection. Actin polymerization was inhibited by 1 μg/ml of cytochalasin D or 250 ng/ml of latrunculin A. Microtubule polymerization was inhibited by 10 μg/ml of colchicine. Stock solutions were dissolved in DMSO unless otherwise indicated and stored at − 20°C. Stock solutions of cytochalasin D, latrunculin A, colchicine, dynasore, nystatin and wortmannin were at 1 mg/ml, 100 μg/ml, 10 mg/ml, 25 mg/ml, 25 mg/ml and 10 mg/ml respectively. Addition of pharmacological inhibitors at the concentrations indicated had no effect on keratinocyte viability, where cytotoxicity assays using the alamarBlue™ cell viability reagent (Invitrogen; Thermo Fisher Scientific, Singapore) was similar to the untreated control (data not shown). Addition of pharmacological inhibitors at the concentrations indicated also had no effect on bacteria viability, where growth kinetics and CFU count were similar to the untreated bacteria control. All chemical inhibitors were purchased from Sigma-Aldrich (St. Louis, MO), unless otherwise stated.

### Immunofluorescence

Following infection, coverslips with cells were washed 3 times in PBS and fixed with 4% paraformaldehyde at 4°C for 15 min. Cells were then permeabilized with 0.1% Triton X–100 (Sigma-Aldrich, St. Louis, MO) (actin) or 0.1% saponin (endosomal compartments) for 15 min at room temperature and washed 3 times in PBS or PBS with 0.1% saponin, respectively. Cells were then blocked with PBS supplemented with 0.1% saponin and 2% BSA. For actin labelling, the phalloidin–Alexa Fluor 568 conjugate (Thermo Fisher Scientific, Singapore) was diluted 1:40 in PBS. For antibody labelling of endosomal compartments, antibody solutions were diluted in PBS with 0.1% saponin at a 1:10 dilution for mouse α-LAMP-1 (ab25630, Abcam, Cambridge, UK), 1:50 for rabbit α-EEA1-Alexa Fluor 647 (ab196186, Abcam, Cambridge, UK), 1:100 for rabbit α-M6PR-568 (ab202535, Abcam, Cambridge, UK) or a 1:100 for rabbit α-cathepsin D (ab75852, Abcam, Cambridge, UK) and incubated overnight at 4°C. The following day, coverslips were washed 3 times in 1X PBS with 0.1% saponin and incubated with a 1:500 dilution of the following secondary antibodies (Thermo Fisher Scientific, Singapore): goat α-Mouse IgG (H+L) Alexa Fluor Plus 647, goat α-Rabbit IgG (H+L) Alexa Fluor Plus 647, goat anti-Rabbit IgG (H+L) Alexa Fluor 568, goat anti-Mouse IgG (H+L) Alexa Fluor 568 for 1 h at room temperature. Coverslips were then washed 3 times in 1X PBS with 0.1% saponin and incubated with a 1:500 dilution of Hoechst 33342 (Thermo Fisher Scientific, Singapore) for 20 min at room temperature. Coverslips were then subjected to a final wash, 3 times with PBS with 0.1% saponin and 2 times with PBS. Coverslips were then mounted with SlowFade™ Diamond Antifade (Thermo Fisher Scientific, Singapore) and sealed.

### Confocal Laser Scanning Microscopy (CLMS)

Confocal images were then acquired with a Zeiss Elyra PS.1 inverted laser scanning confocal microscope (Carl Zeiss, Göttingen, Germany) equipped with three Helium/Neon-lasers (633 nm, 594 nm and 561 nm), one Argon-ion laser (458-514 nm) and one Diode laser (405 nm) using the Zeiss Zen Black 2012 SP2 software suite. Laser power and gain were kept constant between experiments. Control primary and secondary antibody alone labeling experiments were performed in parallel infected cells. Z-stacked images were processed using Zen 2.1 (Carl Zeiss, Göttingen, Germany).

### Construction of LAMP1-mCherry strain

pLAMP1-mCherry (Addgene plasmid #45147, Addgene, Cambridge) and EF1α-mCherry-N1 plasmid (Thermo Fisher Scientific, Singapore) were isolated using the Monarch® Plasmid Miniprep Kit (New England BioLabs Inc., USA), according to manufacturer’s instructions. The LAMP1 gene was then sub cloned into the pEF1α-mCherry-N1 vector using the In-Fusion® HD Cloning Kit (Clontech, Takara, Japan), according to manufacturer’s instructions. The plasmid construct was then transformed into DH5α Stellar™ competent cells by incubating at 42°C for 1 minute. Transformed colonies with the desired construct and primers used for the cloning and subsequent verification are shown in **Supplementary Table 1** with colony PCR. EF1α Lamp1-mCherry plasmid was extracted from successful transformants with the Monarch® Plasmid Miniprep Kit (New England BioLabs Inc., USA), according to manufacturer’s instructions. Keratinocytes were grown in 6-well tissue culture plates as described above, where each well was seeded with 2 ×10^5^ cells. 2.5 μg of plasmid DNA was transfected into keratinocytes using Lipofectamine 3000 (Invitrogen; Thermo Fisher Scientific, Singapore), according to manufacturer’s instructions. The culture media was replaced after 6 h of incubation, followed by a subsequent replacement 18 h later. Keratinocytes were then subjected to Geneticin selection (1 mg/ml) (Invitrogen; Thermo Fisher Scientific, Singapore) to select for transfected clones. Clones stably overexpressing Lamp1-mCherry were subjected to validation by immunoblotting and flow cytometry. Clonal populations were selected and subjected to fluorescence activated cell sorting (FACS) to ensure that the entire population were expressing the fluorescent construct.

**Supplementary Table 1.**
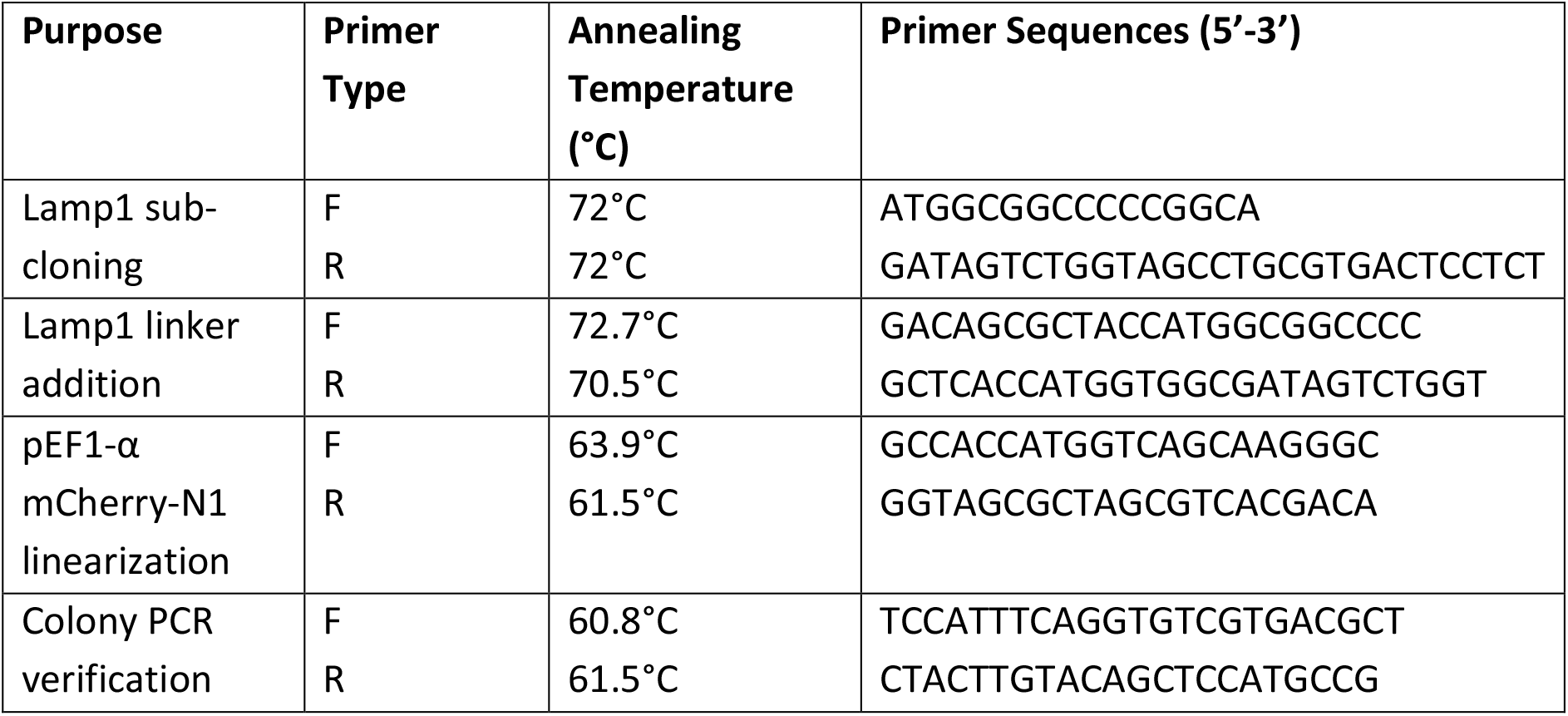
Primers used in this study.

### Correlative light and electron microscopy

*E. faecalis* infected keratinocytes stably transfected with LAMP1-mCherry were grown in 35 mm glass bottom dishes (MatTec Corp., Ashland, USA). Cells were fixed at 18 hpi for 2 h on ice in 2.5% glutaraldehyde (EMS) in 0.1 M cacodylate buffer (CB; EMS) pH 7.4 supplemented with 2 mM CaCl_2_. Cells were washed several times in CB and imaged using a spinning disk confocal microscope (CorrSight, Thermo Fisher Scientific, Singapore). Confocal z-stacks were acquired with a 63x oil objective (NA 1.4, Plan Apochromat M27, Zeiss) on an Orca R2 CCD camera (Hamamatsu, Japan) using standard filter sets. Cells were then further processed for TEM. Briefly, cells were post-fixed with 1% osmium tetroxide for 1 h on ice, washed several times, and incubated with 1% low molecular weight tannic acid ((C14H10O9)n; EMS) for 1 h at RT. Cells were dehydrated using a graded ethanol series (20%, 50%, 70%, 90%, 100%), and embedded in Durcupan resin (Sigma Aldrich). Areas of interest were sawed out of the dish and sectioned parallel to the glass surface by ultramicrotomy (EM UC7, Leica) using a diamond knife (Diatome). Serial 70-80 nm thin sections were collected on formvar- and carbon-coated copper slot grids (EMS). Electron micrographs were recorded on a Tecnai T12 (Thermo Fisher Scientific) TEM operated at 120 kV using a 4k × 4k Eagle (Thermo Fisher Scientific) CCD camera. TEM and confocal microscopy images were manually overlaid and aligned in Photoshop with minimal warping or stretching. Serial Sections were aligned manually in Photoshop, followed by surface rendering in IMOD.

### Immunoblotting

Whole cell (WC) lysates were prepared by adding 488 μL of RIPA buffer (50 mM Tris-HCl, pH 8.0; 1% Triton X-100; 0.5% Sodium Deoxycholate; 0.1% SDS; 150 mM NaCl) to the wells after intracellular infection assays, cells were scrapped and kept in buffer for 30 min at 4°C. Prior to the addition of 74.5 µL of 1 M DTT and 187.5 µL NuPAGE™ LDS Sample Buffer (4X) (Thermo Fisher Scientific), cells were further mechanically disrupted by passing the lysate through a 26g size needle. Samples were then heated to 95°C for 5 min. 15 µL of cell lysate proteins were then separated by 4-12% (w/v) NuPAGE Bis-Tris and transferred to PVDF membranes. Membranes were incubated with Tris-buffered saline, TBS (50 mM Tris, 150 mM NaCl, pH 7.5) containing 0.1% (v/v) Tween-20 (TBST) and 5% (w/v) BSA for 1 h at room temperature. Membranes were incubated with 1:1000 for mouse α-LAMP-1 (ab25630, Abcam, Cambridge, UK), 1:1000 for rabbit α-EEA1-Alexa Fluor 647 (ab196186, Abcam, Cambridge, UK), 1:1000 for rabbit α-M6PR-568 (ab202535, Abcam, Cambridge, UK), a 1:1000 for rabbit α-cathepsin D (ab75852, Abcam, Cambridge, UK), 1:1000 for rabbit α-Rab5-Alexa Fluor 488 (ab270094, Abcam, Cambridge, UK), 1:1000 for rabbit α-Rab7 (ab137029, Abcam, Cambridge, UK), or 1:1000 for rabbit α-GADPH (5174, Cell Signaling Technology) in TBST containing 1% (w/v) BSA overnight at 4°C. Membranes were washed for 60 min with TBST at room temperature and then incubated for 2 h at room temperature with goat anti-rabbit (H+L) or goat anti-mouse HRP-linked secondary antibodies (Invitrogen) respectively. Membranes were washed with TBST for 30 min and specific protein bands were detected by chemiluminescence. Band intensities were quantified relatively to the lane’s loading control using ImageJ2.

### Intracellular Reinfection Assay

Infection of keratinocytes was performed in T175 flasks (Nunc; Thermo Fisher Scientific, Singapore) to harvest intracellular bacteria. The infections were performed similarly as described above, except that keratinocytes were infected at MOI 50 for 3 h and subsequently incubated with gentamicin/penicillin G for 1 h. After disruption of keratinocytes, lysates were collected to harvest the intracellular bacteria. Cell lysates were spun down at 100 × g for 1 min to remove debris and the supernatant, which contained the intracellular bacteria, was transferred into a new tube. Harvested bacteria were washed once in PBS and resuspended in complete DMEM. An aliquot of the bacterial suspension was then used for CFU enumeration. The remainder of the bacterial suspension was used for a second round of infection on keratinocyte monolayers in 6-well plates. To achieve a sufficient MOI with recovered intracellular bacteria, each well of a 6-well plate was infected with intracellular-derived bacteria harvested from a T175 flask. For re-infection studies, keratinocytes were similarly infected with bacteria for 3 h and incubated with gentamicin/penicillin G for 1 h. After disruption of keratinocytes, intracellular bacteria were enumerated and the recovery ratio was determined by calculating the ratio between the inoculum CFU to the recovered intracellular CFU. Parallel infections with planktonically grown bacteria as the inoculum were performed by growing bacteria in complete DMEM for 4 h, before washing once in 0.1% Triton-X100 and a second time in PBS. After resuspending the planktonic bacteria in complete DMEM, bacterial cultures were normalized for infection of keratinocytes at MOI 1, 0.1 and 0.01. For the quantification of total cell-associated bacteria, host cells were infected with intracellular-derived bacteria for 3 h and subsequently lysed for CFU enumeration without prior antibiotic treatment. CFU counts of cell-associated bacteria were normalized against bacterial CFU counts in the supernatant from the same infected wells.

### Statistical analysis

Statistical analysis was done using Prism 9.2.0 (Graphpad, San Diego, CA). We used one- or two-way analysis of variance (ANOVA) with appropriate post tests, as indicated in the figure legend for each figure, to analyze experimental data comprising 3 independent biological replicates, where each data point is typically the average of 3 technical replicates (unless otherwise noted). In all cases, a p value of ≤0.05 was considered statistically significant.

### Ethics statement

All procedures, including isoflurane for anesthesia and CO_2_followed by cervical dislocation for euthanasia, were approved and performed in accordance with the Institutional Animal Care and Use Committee (IACUC) in Nanyang Technological University, School of Biological Sciences (ARF SBS/NIEA-0314).

## Acknowledgements

We thank Kevin B. Wood from the University of Michigan for kindly providing the *E. faecalis* fluorescent reporter plasmid (Dasher) used in this study. We also thank Pei Yi Choo, Sng Wan Xin and Heng Yi Ting Eunice for technical assistance in this project. We are grateful to Kline lab members Haris Antypas, Frederick Reinhart Tanoto, Claudia Stocks, Chor Ming Thong, and Brenda Tien for critical feedback on the manuscript.

Funding for this work was provided by the National Research Foundation and Ministry of Education Singapore under its Research Centre of Excellence Program, by the National Research Foundation under its Singapore NRF Fellowship program to KAK (NRF-NRFF2011-11), by the Ministry of Education Singapore under its Tier 2 programs to KAK (MOE2014-T2-1–129 and MOE2018-T2-1-127), by the Ministry of Education Singapore Tier 1 grants to A.L. (MOE RG136/17 and MOE RG39/14), and by an NTU Start-up grant to A.L. RAGDS is supported by the National Research Foundation, Prime Minister’s Office, Singapore, under its Campus for Research Excellence and Technological Enterprise (CREATE) program, through core funding of the Singapore-MIT Alliance for Research and Technology (SMART) Antimicrobial Resistance Interdisciplinary Research Group (AMR IRG). The funders had no role in study design, data collection and analysis, decision to publish, or preparation of the manuscript.

## Author contributions

WHT and KAK conceptualized the project. WHT, RAGDS, FKH, KKLC, and AL performed experiments and analyzed data. WHT, RAGDS, FKH, AL, and KAK wrote the manuscript; all authors reviewed and approved the final manuscript. AL and KAK provided supervision and acquired funding for the project.

